# Slow integrin-dependent migration organizes networks of tissue-resident mast cells

**DOI:** 10.1101/2022.07.19.500614

**Authors:** Sarah K. Bambach, Lukas Kaltenbach, Nadim Aizarani, Paloma Martzloff, Alina Gavrilov, Katharina M. Glaser, Roland Thünauer, Michael Mihlan, Manuel Stecher, Aude Thiriot, Stephan Wienert, Ulrich von Andrian, Marc Schmidt-Supprian, Claus Nerlov, Frederick Klauschen, Axel Roers, Marc Bajénoff, Dominic Grün, Tim Lämmermann

## Abstract

Many leukocytes use fast and flexible amoeboid migration strategies to move autonomously throughout tissues. Here, we show that the movement of mast cells (MCs), leukocytes with important roles during allergies and anaphylaxis, fundamentally differs from this rapid adhesion-free leukocyte migration. We identify a crucial role for integrin-dependent adhesion in controlling slow MC movement, which shapes the positioning and network-like tissue distribution of this long-lived immune cell type. In contrast to other immune and non-immune cells, MCs cannot compensate for the lack of integrin function by switching to another migration mode. Single-cell RNA-sequencing revealed a special role for integrins in defining a mature MC phenotype in the periarteriolar tissue space where several stromal cell types provide an anatomical niche rich in Kit ligand, the major MC growth and survival factor. Collectively, this study highlights substrate-dependent haptokinesis as an important mechanism for MC network formation and the tissue organization of resident immune cells.

## INTRODUCTION

Coordinated movement, positioning and interaction of immune cells with diverse effector functions build the basis for protective immune responses in our tissues. Leukocyte locomotion through tissues is generally associated with a flexible mode of movement, termed amoeboid migration, which depends on rapid cycles of actin polymerization and actomyosin contraction ^1^. To transmit these intracellular forces to the substrate and move productively, many immune cell types do not necessarily require integrin receptors, the major family of adhesion receptors in mammals ^2^. Several leukocyte types, including neutrophils, T cells and dendritic cells, were shown to move without functional integrins in complex tissue geometries ^3–5^. Numerous recent studies, mostly *in vitro*, revealed that fast amoeboid leukocyte migration can result from a continuum of strategies for force transmission, ranging from very weak integrin-dependent adhesiveness to the utilization of surface topography and unspecific transmembrane coupling ^6–8^. This explains the migratory plasticity of these cells and how they can switch between distinct mechanical modes of migration in order to flexibly adapt to changing external environments ^4^. Together, these studies have led to the generally prevailing paradigm that leukocyte interstitial migration is low– to non–adhesive, which makes immune cells independent of the biochemical composition of the environment and allows them to move autonomously within tissues ^9^. However, whether these generalized concepts apply to the 3D migration strategies of all immune cell types is completely unclear. In particular, it is still largely unknown whether and how integrin-mediated force coupling impacts the movement, positioning and tissue organization of tissue-resident immune cell types.

Mast cells (MCs) are long-lived immune cells that distribute as resident cellular networks throughout tissues that interact with the external environment, including skin, airways and intestine. MCs are best known for their roles in allergy and anaphylaxis, but are also critically involved in the degradation of toxins and immunity against various types of pathogens ^10^. To exert their effector functions, activated MCs release pre-formed inflammatory mediators and proteases ^11^. Moreover, they synthesize substances, whose release can initiate local inflammatory reactions including post-capillary venule dilatation, activation of the blood endothelium, increased blood vessel permeability and recruitment of blood-derived leukocytes ^12^. MCs in the skin connective tissue of adult mice, i.e. mice older than 6 weeks, originate from definitive hematopoiesis and maintain themselves independently from the bone marrow ^13–15^. Their differentiation and survival critically depend on Kit ligand (KITLG, also known as stem cell factor, SCF), which signals through the receptor tyrosine kinase KIT (c-KIT) ^16^. How MCs move and strategically position to maintain their survival and cellular network distribution in physiological tissues is completely unknown. Intravital imaging studies revealed extremely slow, if any, MC kinetics over several hours ^17, 18^. MCs are largely found within the fibrillar interstitium, but also locate closely to blood vessels and nerves ^19–21^. However, the molecular mechanisms that underlie such site-specific MC localization have not been investigated. All current concepts on MC migration rely on *in vitro* studies, while the mechanisms controlling the cytoskeletal dynamics of MC movement in tissues have never been addressed. Although murine and human MCs are known to express integrin receptor classes that could interact with other cells or extracellular matrix (ECM) ^22–24^, it has remained unanswered whether and how these adhesion receptors control MC migration, positioning and their organization in the native tissue environment.

## RESULTS

### Mast cells form integrin-dependent adhesion structures to interact with ECM

To examine the positioning of endogenous MCs in real tissue, we used *Mcpt5-Cre^+/−^ Rosa26^LSL:Tom^* mice to visualize connective-tissue type MCs (CTMCs) in relation to ECM structures of the dermal skin. TdTomato-expressing cells were identified as avidin-positive dermal MCs, which were either aligned along vascular basement membranes or surrounded by fibrillar fibronectin (FN) in the interstitial space (Fig. 1a,b). To investigate the nature of MC-ECM interactions, we generated bone marrow-derived mast cells (BMMCs) that were differentiated toward a CTMC phenotype ^25^. These primary MCs highly expressed ECM-binding integrin heterodimers of the β1 and αv family (ED Fig. 1). When placed on fibroblast-derived FN matrices, BMMCs formed integrin α5β1-containing adhesion structures reminiscent of fibrillar adhesions in fibroblasts (Fig. 1c,d) ^26^. BMMCs also spread on FN-coated 2D surfaces in the presence of KITLG (Fig. 1e) or upon FcεR1 engagement (Fig. 1f) ^27, 28^. Under both conditions, BMMCs formed focal adhesion structures, which were detected based on the presence of integrin α5β1, paxillin, talin, vinculin and activated β1 integrin (9EG7) predominantly at the cell periphery (Fig. 1e,f). Thus, our results demonstrate prominent integrin-containing adhesion structures in MCs interacting with several forms of ECM.

**Fig. 1.**
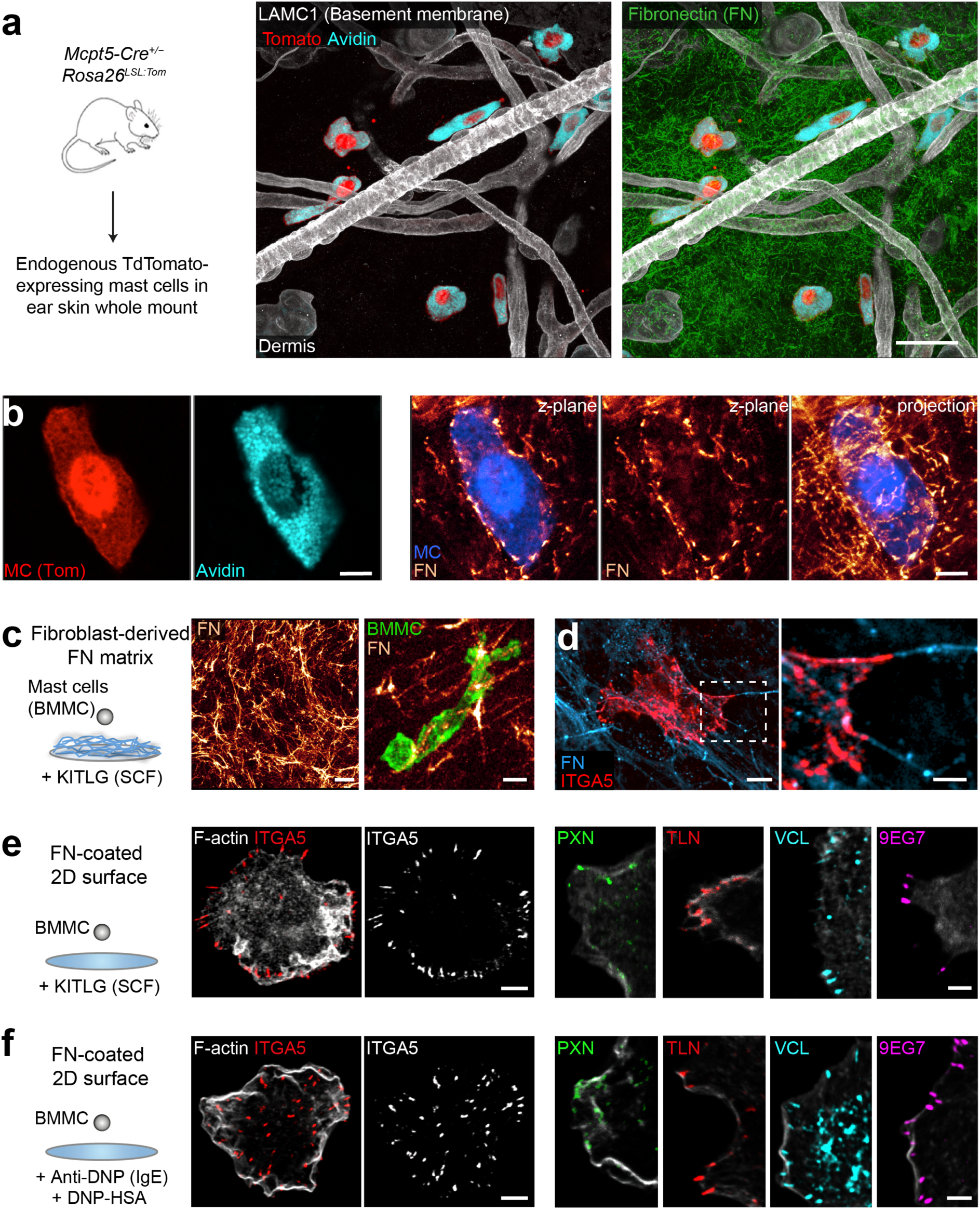
Mast cells form integrin-based adhesions to interact with ECM. **(a, b)** Immunofluorescence staining of ear skin whole mount tissue from an adult *Mcpt5-Cre^+/−^ Rosa26^LSL:Tom^* mouse. (a) Overview image of Tomato-expressing and avidin-positive endogenous mast cells (MCs) (red, cyan) in relation to LAMC1-positive basement membrane components (white) and the interstitial matrix protein fibronectin (green) in the skin dermis. (b) High magnification image of a Tomato- and avidin-positive interstitial MC (left), which is enwrapped by a fine network of fibronectin (FN) fibers of the native dermis (right). Single *z*-planes and a 3-µm-projection are shown. FN fibers are displayed as glow heatmap. **(c)** Interaction of living Lifeact-GFP expressing bone marrow-derived MCs (BMMCs, green) with a fluorescent fibroblast-derived FN fibril network *in vitro*. **(d)** Confocal fluorescence microscopy of a BMMC fixed and immunostained for integrin α5 (ITGA5) (red). Zoom-in image (right) shows adhesion structures along FN fibers. **(e, f)** Interaction of BMMCs with FN-coated glass slides in the presence of Kit ligand (KITLG) (e) or 30 min after IgE/DNP-HSA-induced cell spreading (f). Confocal fluorescence microscopy of BMMCs stained for F-actin and the focal adhesion components integrin α5, paxillin (PXN), pan-talin (TLN), vinculin (VCL) and active integrin β1 (9EG7). Scale bars: 30 µm (a, d left), 15 µm (d right), 5 µm (b, c, e left, f left), and 2 µm (e right, f right). See also Extended Data Fig. 1.

### MC migration requires high-affinity integrins on 2D surfaces and in confined spaces

To interfere with integrin functionality, we generated *Mcpt5-Cre^+/−^ Tln1^fl/fl^* mice to deplete talin- 1 in BMMCs (Fig. 2a and ED Fig. 2a). Talin-1 interacts with integrin cytoplasmic domains and is crucial for integrin activation, ligand binding and coupling of F-actin to adhesion sites ^29^. As hematopoietic cells express only low levels of the talin-2 isoform, TLN1 depletion efficiently reduced pan-talin protein levels in BMMCs (ED Fig. 2b,c) without altering MC maturation and integrin cell surface expression (ED Fig. 2d,e). As expected *Tln1^−/−^* BMMCs did not bind to ECM substrates, which WT BMMCs adhered to under several experimental conditions (Fig. 2b). Live cell imaging (Fig. 2c and Supplementary Video 1) and 2D spreading analysis (Fig. 2d,e) corroborated this adhesion deficit. Similarly impaired adhesion responses were observed for *Tln1^−/−^* dermal MCs (DMCs) and peritoneal MCs (PMCs) (ED Fig. 2a,f). Consequently, *Tln1^−/−^*BMMCs could not invade fibroblast-derived FN matrices (ED Fig. 2g and Supplementary Video 2).

**Fig. 2.**
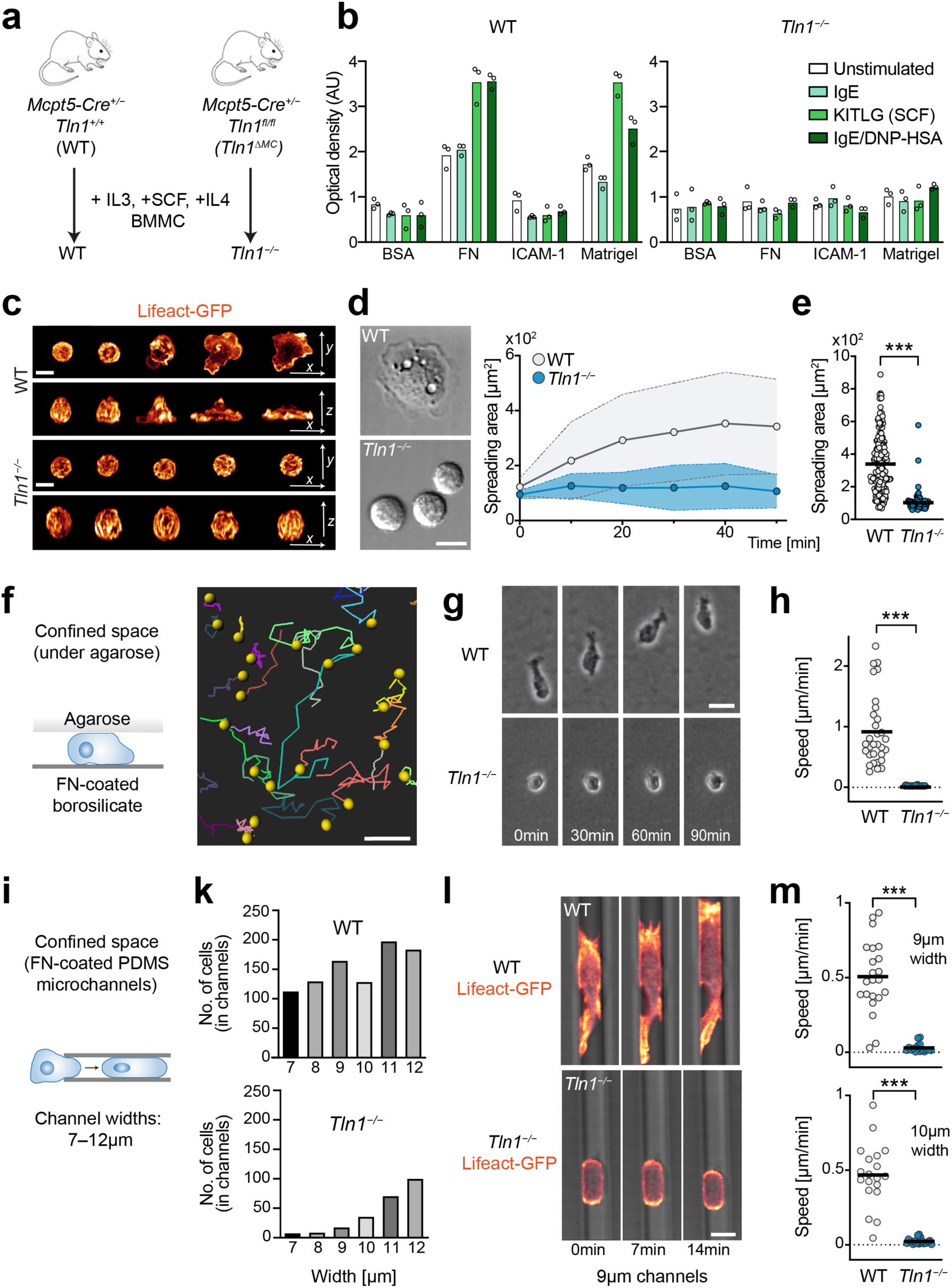
High-affinity integrins are critical for MC migration on 2D surfaces and in confined spaces. **(a)** Scheme for the generation of BMMCs depleted of talin-1. **(b)** Adhesion assay of WT and *Tln1^−/−^* BMMCs to several integrin ligands upon stimulation with IgE, KITLG or IgE/DNP-HSA. Bovine serum albumin (BSA) assessed background adhesion. Bar charts represent the average optical density (OD) value of adherent cells. Data from one representative experiment (*N*=3 technical replicates, mean±SD) of *n*=3 independent experiments for each condition are shown. **(c–e)** MC spreading on FN-coated 2D surfaces in response to IgE/DNP-HSA. (c) Spinning disk-confocal microscopy of Lifeact-GFP expressing WT and *Tln1^−/−^*BMMCs. A time sequence from live cell microscopy over 20 min is displayed. Confocal *z*-stacks are shown as merged projection (*x-y*-plane: upper row, x-*z*-plane: lower row). (d) Graphical analysis of the MC spreading area over time, and representative differential interference contrast (DIC) images of WT and *Tln1^−/−^*BMMCs at 30 min. Dots in the graph display mean±SD values from one representative experiment. (e) Comparative analysis of WT and *Tln1^−/−^* BMMC spreading area after one hour. Dots represent values of individual cells from one representative experiment of *n*=4 independent experiments for each genotype. Bars display the mean; \*\*\**P*<0.001, *U* test. **(f–h)** BMMC migration in the confined space of an under-agarose assay was recorded with live cell imaging. (f) Trajectories of individual WT cell tracks over 14 hours are shown. (g) Time-lapse sequences over 90 min of WT and *Tln1^−/−^*BMMC morphologies and movement. (h) Quantification of the average cell speed from one representative experiment of *n*=3 independent experiments for each genotype are shown. Dots are values of individual cells (*N*=20–30 randomly chosen cells per genotype). Bars display the mean; \*\*\**P*<0.001, *t* test. **(i–m)** BMMC migration in the confined space of PDMS microchannels was analyzed (i). (k) WT and *Tln1^−/−^* BMMC invasion into channels of varying width was quantified. Data from one multichannel experiment per genotype are displayed. (l) Spinning disk-confocal microscopy of Lifeact-GFP expressing WT and *Tln1^−/−^* BMMCs migrating in 9-µm-wide channels. A time sequence from live cell microscopy over 14 min is displayed. Lifeact-GFP expression is displayed as glow heatmap. (m) Quantification of the average cell speed in channels of 9 µm and 10 µm width from one multichannel experiment per genotype are shown. Dots are values of individual cells (*N*=16–22 cells per condition). Bars display the mean; \*\*\**P*<0.001, *U* test. Scale bars: 10 µm (c, d, l), 30 µm (g), and 100 µm (f). See also Extended Data Fig. 2 and Supplementary Videos 1–5.

Next, we tested the migration potential of *Tln1^−/−^* BMMCs in confined spaces, which allow specific cell types to switch from adhesion-dependent to adhesion-free migration modes ^30, 31^. First, we examined MCs moving in an under-agarose assay setup (Fig. 2f), which had previously been shown to support the rapid movement of integrin-deficient dendritic cells ^31^. Live imaging over 14 hours revealed that WT BMMCs perform lamellipodia-based migration at median speeds of < 1 µm/min, which depended on F-actin dynamics (ED Fig. 2h) and actomyosin contraction (ED Fig. 2i). In contrast, *Tln1^−/−^* BMMCs formed only rudimentary cell protrusions and remained largely immotile (Fig. 2g,h and Supplementary Video 3). Total interference reflection (TIRF) microscopy of Lifeact-GFP dynamics demonstrated the insufficient coupling of actin flow to the surface, by showing a clear increase in the retrograde actin flow of *Tln1^−/−^* BMMCs (ED Fig. 2k and Supplementary Video 4). Second, we studied MC migration inside FN-coated microchannels of varying width (Fig. 2i). Not only did *Tln1*- deficiency impair MC invasion into the channels (Fig. 2k), but also stalled the movement of cells that had managed to enter the confined space (Fig. 2l,m and Supplementary Video 5). Thus, our data show that integrin-dependent force coupling to ECM is crucial for MC adhesion and movement on 2D surfaces, cell-derived matrices and in confined environments.

### Integrins control MC migration, positioning and periarteriolar alignment in real tissue

Ultimately, we were interested in understanding MC interstitial movement in real tissues. Physiological tissue environments are extremely heterogeneous, providing migrating immune cells many options in forms of 2D surfaces, confined spaces and 3D networks in close proximity to one another ^3^. We chose to analyze the homeostatic ear skin of adult mice, where the residing endogenous MCs organize as networks of homogenously distributed individual cells throughout the connective tissue of the dermis. To interfere with integrin functionality in these cells, we used *Mcpt5-Cre^+/−^ Tln1^fl/fl^*(short: *Tln1^ΔMC^*) mice with specific conditional deletion of talin-1 in CTMCs. Comparison of MC distribution in ear skin whole mount preparations of *Tln1^ΔMC^*and control animals revealed two striking phenotypes. First, *Tln1*-deficient MCs lost their homogenous tissue distribution and instead formed cellular clusters in the dermis (Fig. 3a,b). This phenotype was not observed in *Rag2^−/−^* mice, ruling out a role of immunoglobulin E (IgE)-stimulated MC adhesion in this process (ED Fig. 3a) ^28^. Dermal MC numbers were comparable between *Tln1^ΔMC^* and control mice (ED Fig. 3b). Second, *Tln1*-deficient MCs lost their localization along dermal arterioles (Figure 3b). By using two experimental strategies to identify dermal vessel types, we show that the perivascular alignment in *Tln1^ΔMC^*mice is only affected at arterioles, but not at postcapillary venules or capillaries (Fig. 3b,c and ED Fig. 3c,d). Consequently, MC coverage of the outer arteriolar walls was diminished in knockout animals (Fig. 3d). Cell shape analysis of endogenous, YFP-expressing CTMCs revealed obvious differences: WT MCs displayed elongated, mesenchymal-like morphologies with lamellopodial-like leading edges, whereas *Tln1*-deficient MCs showed round, amoeboid-like shapes (Fig. 3e,f). Given the important roles of β1 integrins for cellular mechanotransduction in many non-immune cells ^32^, we also analyzed *Mcpt5-Cre^+/−^ Itgb1^fl/fl^* (*Itgb1^ΔMC^*) mice with CTMC-specific depletion of the *Itgb1* gene. These animals phenocopied *Tln1^ΔMC^* mice with similar decreases in the percentage of periarteriolar MC and arteriolar MC coverage (Fig. 3g-i). Moreover, *Itgb1*-deficient MCs also showed a mesenchymal-to-amoeboid shape switch (Fig. 3k, ED Fig. 3e). These phenotypes were not observed in *Itgb2^−/−^* and *Vav-iCre Itgb3^fl/fl^*mice (data not shown). Thus, our results demonstrate a crucial role of ECM-binding integrins for maintaining MC shape, positioning and organization in living connective tissue.

**Fig. 3.**
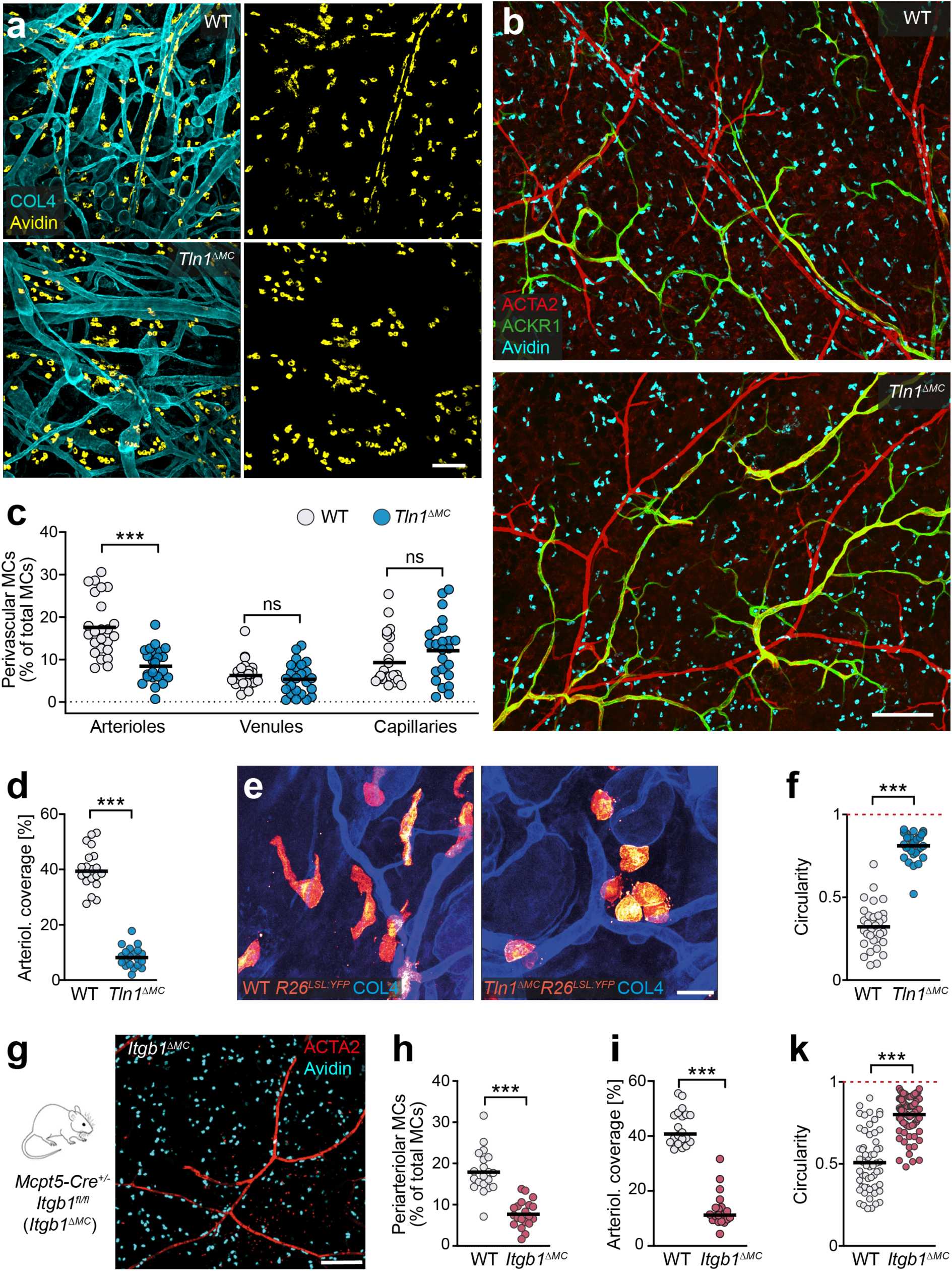
Integrins control MC network formation and periarteriolar alignment *in vivo*. **(a, b)** Comparative analysis of ear skin whole mount tissues of adult *Mcpt5-Cre^+/−^ Tln1^fl/fl^* (*Tln1^ΔMC^*) mice and littermate controls. Endogenous dermal MCs were immuno-stained with avidin in relation to collagen IV (COL4)- expressing basement membrane structures (a) or dermal vessel types (b). Arterioles (red) and postcapillary venules (green) were classified by differential staining for α-smooth muscle actin (ACTA2) and ACKR1. **(c, d)** Analysis of perivascular MC positioning. (c) Quantification of MCs in proximity to arterioles, venules and capillaries, which were identified by differential staining for ACTA2 and endomucin. (d) Quantification of arteriolar coverage by MCs. For both (c) and (d), dots represent individual imaging fields of view, which come from *n*=6 (c) and *n*=5 (d) 7-week old mice; \*\*\**P*<0.001; NS, non-significant; two-way ANOVA (c) and *t* test (d). **(e, f)** Analysis of MC morphologies in the dermis of adult *Mcpt5-Cre^+/−^ Tln1^fl/fl^ Rosa26^LSL:YFP^* mice and littermate control *Mcpt5- Cre^+/−^ Tln1^+/+^ Rosa26^LSL:YFP^*mice. (e) YFP-expressing dermal MCs are displayed in glow heatmap. (f) Quantification of cell roundness (circularity) was performed for *N*=31 cells per genotype, which came from *n*=4 (WT) and *n*=3 (*Tln1^ΔMC^*) mice. Bars display the mean; \*\*\**P*<0.001, *U* test. **(g–k)** Analysis of dermal MCs in 7- to 10-week old *Mcpt5-Cre^+/−^ Itgb1^fl/fl^* (*Itgb1^ΔMC^*) and littermate control mice: MC distribution in ear skin whole mounts (g), and quantification of arteriolar MC numbers (h), arteriolar coverage (i) and circularity (k) in comparison to littermate control mice. For both (h) and (i), dots represent individual imaging fields of view, which come from *n*=5 mice per genotype; \*\*\**P*<0.001, *t* test. (k) Quantification of cell roundness (circularity) was performed for *N*=59 (WT) and *N*=91 (*Itgb1^ΔMC^*) cells, which came from *n*=3 mice per genotype. Bars display the mean; \*\*\**P*<0.001, *U* test. Scale bars: 100 µm (a), 200 µm (b, g), and 20 µm (e). See also Extended Data Fig. 3 and Materials and Methods section.

### MCs use slow integrin-dependent migration in tissues and fibrillar matrices

Dermal MCs are long-lived cells with slow proliferation rates, maintaining themselves during homeostasis in adult mice ^13–15^. They are considered sessile cells with very slow migration dynamics ^17, 18^. We hypothesized that the observed MC clusters in *Tln1^ΔMC^* mice resulted from an MC migration deficit (Fig. 3a,b). We speculated that TLN1 knockout MCs could still proliferate at slow rate, but then failed to move and invade the surrounding interstitial space. To address this question directly in the tissue, we crossed *Tln1^ΔMC^* mice with transgenic Ubow mice, which have previously been used for *in situ* fate tracking of other immune cell types ^33^. In these mouse crosses, *Mcpt5* promoter activity drives a Cre recombinase-mediated single and definitive recombination event, by which any Ubow MC acquires a specific color, either YFP or CFP at ∼2:1 stochastics, that is stable for the life of the cell and inherited to all its progeny (Fig. 4a). Analysis of ear skin whole mounts of control Ubow mice revealed a heterogenous mixed distribution of YFP^+^ and CFP^+^ MCs in the adult skin dermis (Fig. 4b). In contrast, *Tln1^ΔMC^ Ubow* mice showed areas of unicolored MC clusters, strongly suggesting that MCs originated from a few parent MCs in these regions, but then did not move further into the interstitial dermal space to intermix with other MCs (Fig. 4b). Detailed quantification of cell cluster size with ClusterQuant2D software ^33^ showed that WT MCs distributed predominantly as single cells or two-cell pairs, which we did not categorize as cell cluster, whereas *Tln1*-deficient MCs were commonly associated to clusters of > 3 cells (Fig. 4c and ED Fig. 4a,b).

**Fig. 4.**
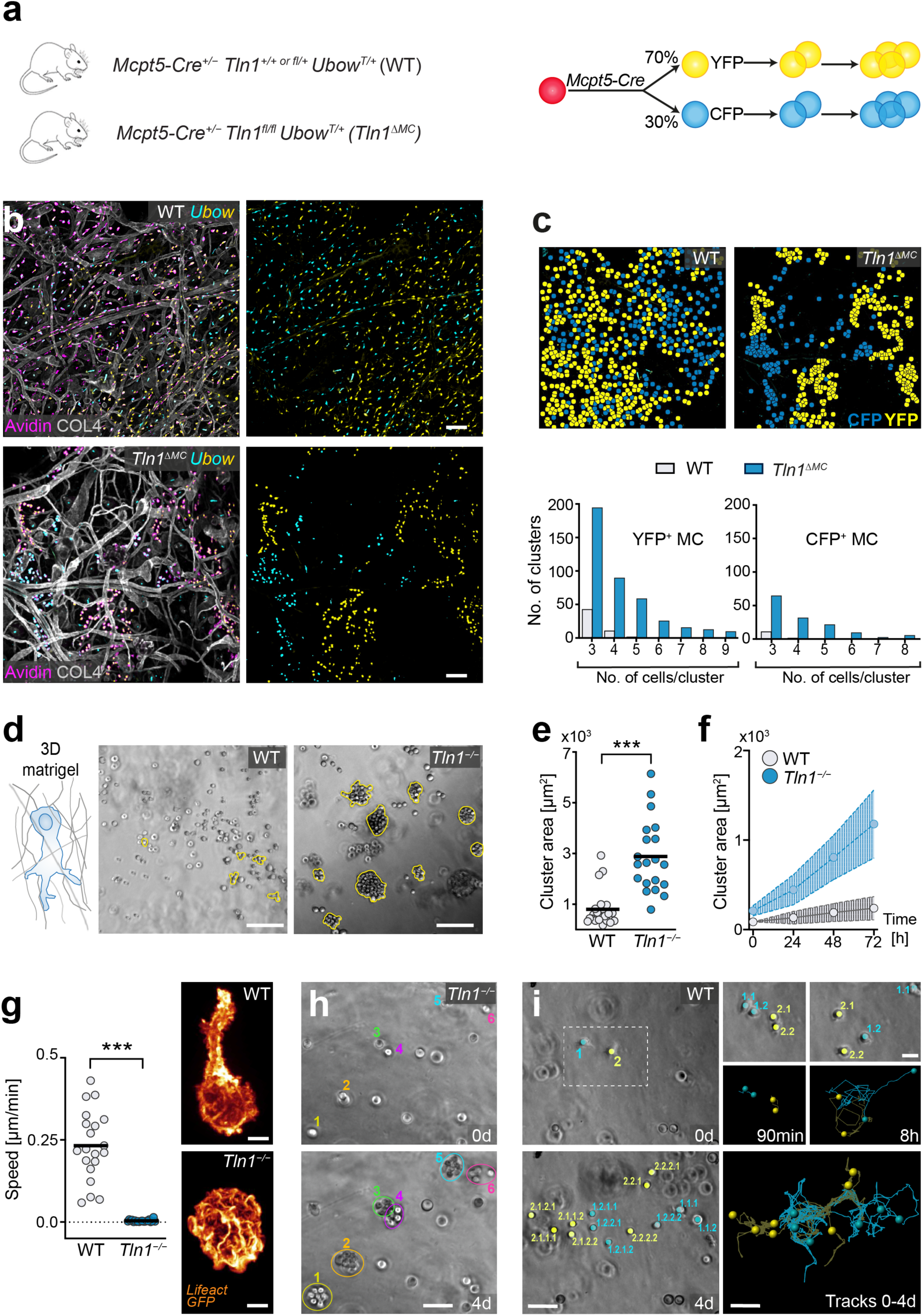
MCs use slow integrin-dependent migration in tissues and fibrillar matrices. **(a)** Experimental strategy for tracing slow MC migration in the dermal tissue of 6- to 12-week old mice. *Tln1^ΔMC^* and littermate control mice were crossed with Ubow transgenic mice. Cre expression driven by the *Mcpt5* promoter leads to the expression of either fluorescent YFP or CFP in dermal MCs. The descendant cells will carry the same fluorescent label, and the distribution of colors can be followed in adult skin tissue. **(b)** Comparative analysis of ear skin whole mount tissue of 8-week old WT Ubow and *Tln1^ΔMC^* Ubow mice. All YFP- and CFP- positive cells were also positive for the pan-MC marker avidin (purple). Dermal tissue was counterstained with collagen IV (white) to assess basement membranes and overall tissue geometries. **(c)** Representative images of cell center coordinates upon computational rendering using ClusterQuant software, which was used to analyze the formation and size of unicolored MC clusters (see also Extended Data Fig. 4a). Quantification of YFP^+^ and CFP^+^ cluster frequencies sorted by cluster size. Two-cell clusters were not included in the analysis. Data came from *n*=3 (WT) and *n*=4 (*Tln1^ΔMC^*) mice with 2–3 imaging field of views per mouse. **(d–f)** WT and *Tln1^−/−^*BMMCs were placed in 3D fibrillar matrigel and monitored over 3 days. Brightfield images of WT and *Tln1^−/−^* BMMC distribution in gels are displayed (*t*=72 h) (d). (e) Quantification of the cell cluster area (*t*=72h). Each dot represents the growth area of one MC cluster, obtained from *n*=3 independent experiments. Bars display the mean; \*\*\**P*<0.001, *t* test. (f) Analysis of MC cluster growth area over time, dots show individual clusters (mean+/−SD, *N*=10 clusters from one independent experiment). **(g–i)** BMMC migration in 3D matrigel was recorded by live cell microscopy over 4 days. (g) Quantification of the average cell speed is shown. Dots are values of individual cells (*N*=20–25 randomly chosen cells per genotype). Bars display the mean; \*\*\**P*<0.001, *t* test. Spinning-disk confocal images of Lifeact-GFP expressing WT and *Tln1^−/−^* BMMC migrating in matrigel. (h) Time-lapse microscopy revealed proliferation of individual *Tln1^−/−^* BMMC (colored numbers), but hardly any movement out of cell clusters. (i) Live cell imaging of WT BMMCs over 4 days and tracking analysis revealed intermittent phases of movement and cell division, which distributes WT cells in the gel. Colored numbers indicate parent and descendant cells. Scale bars: 100 µm (b, d), 5 µm (g), 50 µm (h, i), and 20 µm (i, insert). See also Extended Data Fig. 4 and Supplementary Videos 6 and 7.

To proof that cluster formation resulted from a migration deficit of adhesion-deficient MCs, we mimicked the dermal interstitial space *in vitro* and imaged BMMC cultures over several days in 3D matrigel (Fig. 4d). WT BMMC moved at median speeds of 0.1–0.3 µm/min in this setting, critically depending on F-actin dynamics (ED Fig. 4c), while actomyosin contraction had only a contributing role (ED Fig. 4d). Comparison of WT and *Tln1^−/−^* BMMC cultures over 4 days showed a clear difference that matched our *in situ* observations: WT cells distributed throughout the gel, whereas *Tln1^−/−^* cells formed MC clusters over time (Fig. 4d–f). Live cell imaging and cell tracking showed that the lack of integrin functionality caused BMMCs to adopt round, amoeboid-like shapes, which did not support migration in 3D gels (Fig. 4g). As a consequence, locally proliferating *Tln1^−/−^*BMMCs underwent cell division, but then did not move into the surrounding gel, resulting in MC cluster formation (Fig. 4h and Supplementary Video 6). Only in very rare cases we observed amoeboid-like migrating *Tln1^−/−^* BMMCs, which, if it happened, could seed of a new MC cluster (ED Fig. 4e and Supplementary Video 7). In contrast, WT BMMCs moved away from each other after cell division, allowing a few individual parent cells to completely seed interstitial gel matrix over 4 days (Fig. 4i and Supplementary Video 6). Together, we demonstrate the MCs strictly depend on integrin-mediated force coupling to ECM for productive migration in artificial and physiological 3D environments. This mode of movement allows MC organization as homogenously distributed individual cells throughout interstitial spaces.

### Single-cell RNA-seq analysis reveals a phenotypic subset of dermal MCs

To investigate potential consequences resulting from the altered MC tissue distribution in *Tln1^ΔMC^* mice, we performed single-cell RNA-sequencing (scRNA-seq) of endogenous MCs from WT and knockout mice. We sorted YFP-expressing cells from digested ear skin of *Mcpt5-Cre^+/−^ Rosa26^LSL:YFP^*and *Tln1^ΔMC^ Rosa26^LSL:YFP^* mice, followed by sequencing and computational analysis (Fig. 5a). Based on the expression of the MC marker *Cpa3*, almost all sorted cells were identified as MCs (ED Fig. 5a-c). Only a very small cell subset showed additional macrophage features (cluster 10) (ED Fig. 5c), which we excluded from further analysis. A UMAP representation of single-cell transcriptomes of sorted CD45^+^Lin^−^YFP^+^ cells highlights nine clusters of skin MCs as defined by RaceID3 ^34^ (Fig. 5b). Direct comparison of MCs isolated from WT or *Tln1*^ΔMC^ mice highlighted three clusters (1, 8, 9) enriched for WT cells (Fig. 5c,d). Closer examination of differentially expressed genes in these three clusters revealed two transcriptomic signatures: (1) an upregulation of several MC differentiation and maturation markers, including *Mcpt5 (Cma1)*, *Mcpt6 (Tpsb2)*, *Mcpt4*, *Cpa3* (Fig. 5e,f) ^11^, and (2) a cytoskeletal signature based on the upregulation of transcripts for the intermediate filaments *Vim* and *Lmna* and for the actin regulators *Cdc42* and *Arpc2* (ED Fig. 5d). Thus, integrin-mediated adhesion supports the establishment of a specific MC phenotype in dermal tissue.

**Fig. 5.**
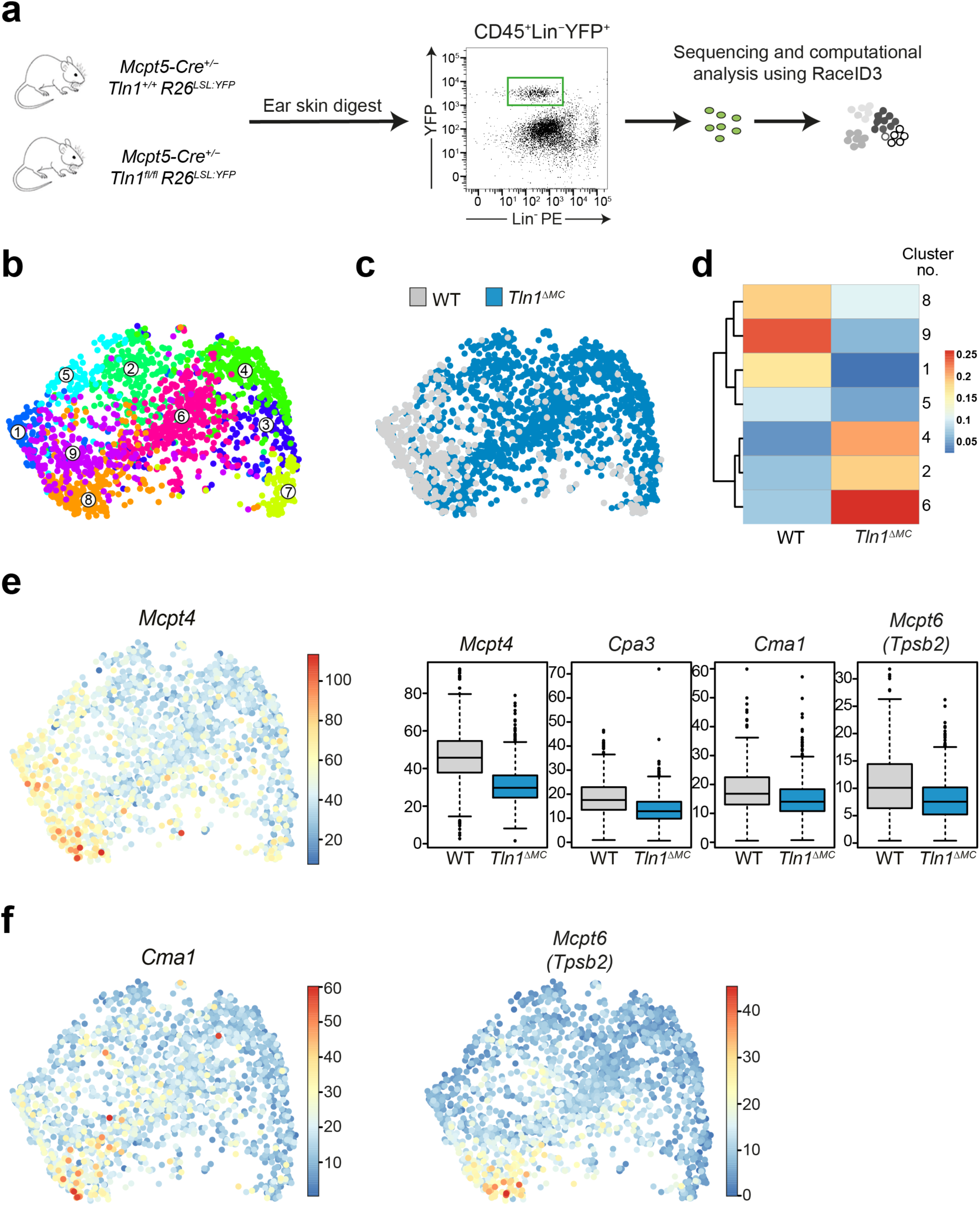
Single cell RNAseq analysis reveals a phenotypic subset of dermal MCs. **(a)** Experimental strategy and work flow for scRNAseq analysis of MCs from dermal ear skin. **(b)** UMAP of single-cell transcriptomes of sorted CD45^+^Lin^−^YFP^+^ cells (WT and *Tln1*^ΔMC^ MCs combined) highlighting RaceID3 clusters. All displayed cells highly express *Cpa3* (see Extended Data Fig. 5b). Numbers denote clusters. **(c)** UMAP of single-cell transcriptomes of MCs highlighting the sources or origins of the cells, i.e. which mice (WT or *Tln1*^ΔMC^) the cells were isolated from. **(d)** Heatmap showing the fractions of WT and *Tln1*^ΔMC^ MCs in clusters, which were significantly enriched for either WT or *Tln1*^ΔMC^ cells;. *P*<0.05, hypergeometric test. **(e)** UMAP showing the expression of *Mcpt4* (left). The color bar indicates normalized expression or transcript counts. Box plots show analysis of clusters 1, 8, 9 and the expression of the MC markers *Mcpt4*, *Cpa3, Cma1, and Mcpt6/Tpsb2* in WT and *Tln1*^ΔMC^ cells (right). The expression of the MC markers was significantly downregulated in MCs from *Tln1*^ΔMC^ relative to WT mice. For all comparisons: Benjamini-Hochberg corrected *P*<0.05 (see Methods). **(f)** UMAP showing the expression of *Cma1 and Mcpt6 (Tpsb2).* The color bar indicates normalized expression or transcript counts. See also Extended Data Fig. 5.

### Integrin-dependent adhesion confines a mature MC subset to arterioles

As phenotypic MC heterogeneity had not been previously described in homeostatic mouse skin, we sought to validate the differential expression of MC marker genes in the intact tissue. To identify MC subsets and relate them to anatomical tissue structures, we established fluorescent *in situ* hybridization (FISH) for *Mcpt5 (Cma1)* and *Mcpt6* in combination with antibody-based immunofluorescence labeling of ear skin whole mounts (Fig. 6a and ED Fig. 6a). *Cma1* and *Mcpt6* FISH signals were increased in MCs aligning along arterioles in comparison to MCs in the interstitial space (Fig. 6b-d). This unexpected finding was confirmed by mMCP-6 protein detection with antibody-based immunofluorescence stainings (ED Fig. 6b-d). While the majority of interstitial MCs showed only intermediate fluorescence signals of mMCP-6 staining, ∼80% of periarteriolar MCs displayed strong mMCP-6 signals (Fig. 6e-g, i). Next, we analyzed mMCP-6 expression in *Tln1^ΔMC^* mice. Of the much lower number of MCs interacting with arterioles in these mice (Fig. 3c), the same percentage of MCs still expressed strong mMCP-6 signals (Fig. 6h,i). In other words, the close vicinity of MCs to arterioles, even in the absence of integrin engagement, is sufficient to promote expression of the MC maturation marker mMCP-6. Thus, our findings determine a crucial role of integrin receptors for anchoring a mature MC phenotype to the periarteriolar compartment.

**Fig. 6.**
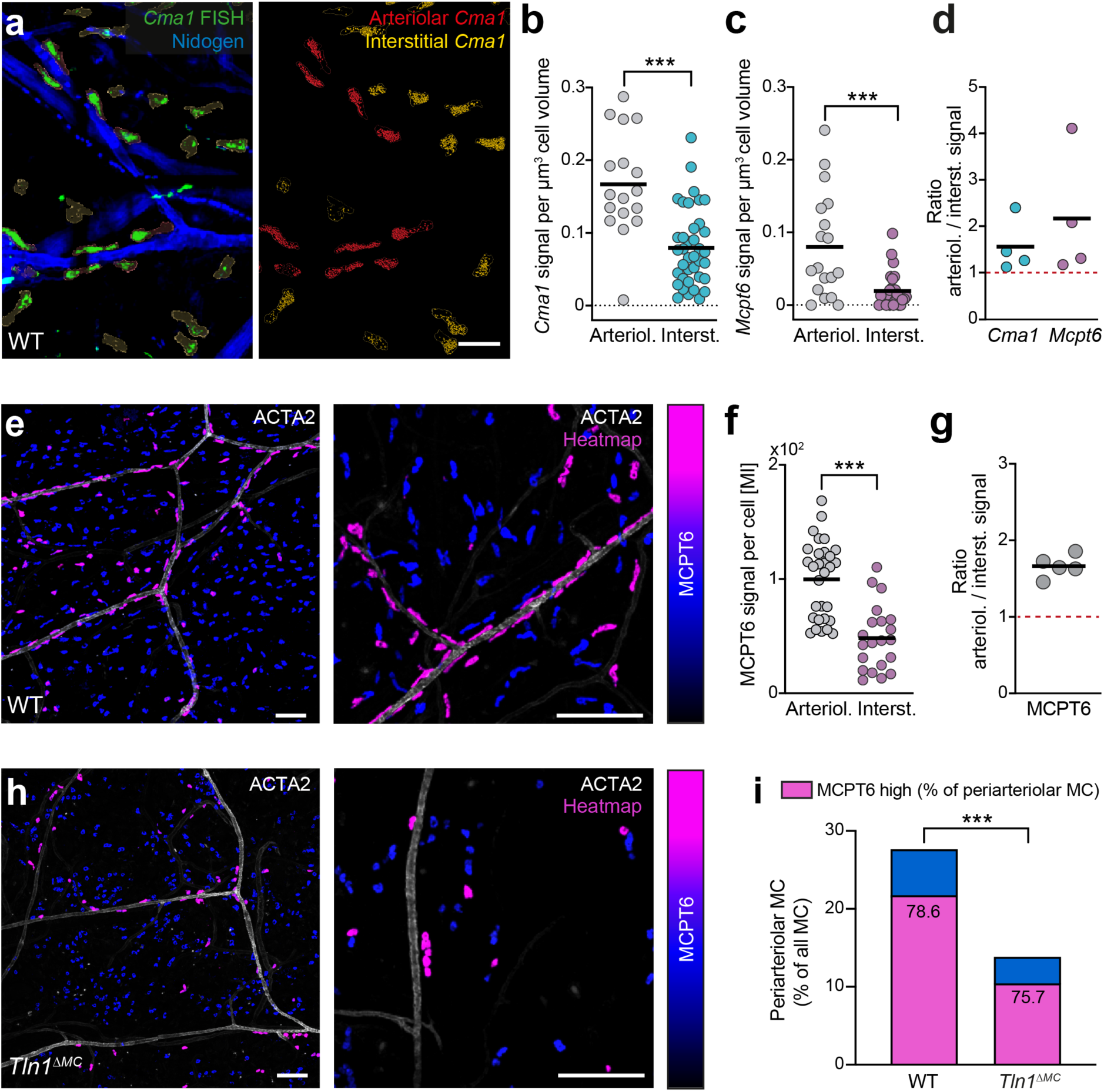
Integrin-dependent adhesion confines a mature MC subset to arterioles. **(a)** Visualization of *Cma1* mRNA using fluorescence in situ hybridization (FISH) in ear skin whole mount of adult WT mice (left) and post-imaging analysis of *Cma1* expression in periarteriolar versus interstitial MCs (right). **(b–d)** Quantification of *Cma1* mRNA (b) and *Mcpt6* mRNA (c) FISH signal per cell volume was performed and compared between periarteriolar versus interstitial MCs. The analysis of one representative imaging field of view is displayed, each dot represents one MC. Bars indicate mean; \*\*\**P*<0.001, *t* test (b, c). (d) Ratios of arteriolar versus interstitial mRNA signals were calculated for independent experiments (*n*=4 mice). **(e–g)** MCPT6 protein expression was analyzed in dermal MCs of adult WT mice. (e) Immunofluorescence staining for MCPT6 in ear skin whole mount tissue: overview image (left), MCs in the periarteriolar space (right). MCPT6 fluorescence intensity is displayed as heatmap. (f) Quantification of MCPT6 fluorescence. The mean intensity (MI) of MCPT6 fluorescence was measured per cell in arteriolar versus interstitial MCs. Quantification is displayed for one representative imaging field of view, each dot represents one MC. Bars indicate mean; \*\*\**P*<0.001, *t* test. (g) Ratios of arteriolar versus interstitial MCPT6 signal were calculated for independent experiments (*n*=5 mice). **(h)** Immunofluorescence staining for MCPT6 in ear skin whole mount tissue of an adult *Tln1^ΔMC^* mouse, data presentation as for WT in (e). **(i)** Comparative analysis of MCPT6 expression in periarteriolar MCs was performed for WT and *Tln1^ΔMC^* mice. The full bar displays the percentage of periarteriolar MCs of total dermal MCs (in analogy to Fig. 3c); \*\*\**P*<0.001, *t* test. Bars also display the percentage of MCPT6^high^ expressing MCs (purple color) within this subset of periarteriolar MCs; *P*>0.05, *t* test. MCPT6^low^ expressing MCs at arterioles are represented in blue. Data includes the analysis of three imaging field of views from *n*=5 (WT) and *n*=4 (*Tln1^ΔMC^*) mice. Scale bars: 20 µm (a), 100 µm (e left, h left), and 150 µm (e right, h right). See also Extended Data Fig. 6.

### KITLG-expressing stromal cells in the periarteriolar space interact with mMCP-6^high^ MCs

Our immunofluorescence analyses suggested that periarteriolar mMCP-6^high^ MCs are in direct contact with the ACTA2^+^ vascular smooth muscle cell (VSMC) layer, which surrounds arteriolar endothelial cells (Fig. 3b, Fig. 6e, ED Fig. 3c,d, ED Fig. 6b–d) ^35^. Hypothesizing that VSMCs might contribute factors that promote a mature MC phenotype, we performed scRNA-seq analysis of GFP-positive cells from digested ear skin of transgenic *Myh11-GFP* transgenic mice. Cells were sorted for GFP^high^ expressing VSMCs, but also collected GFP^low^ pericytes and a small fraction of fibroblasts (ED Fig. 7a–d). Computational analysis of the sequencing data revealed higher expression of Kit ligand (Kitl), the major MC growth, survival and differentiation factor, in VSMCs in comparison to the other two stromal cell types (ED Fig. 7e,f). To confirm this finding directly in the tissue, we analyzed *Kitl* promoter activity in two reporter mouse strains, *Kitl^GFP^* knockin ^36^ and transgenic *Kitl-TdTomato* mice ^37^. ACTA2^+^ arterioles displayed high fluorescent reporter signal in both mouse strains, revealing strong *Kitl* promoter activity as an indication of KITLG expression in this dermal tissue compartment (Fig. 7a,b). Moreover, MCs along arterioles displayed strong fluorescence signals of nuclear localized microphthalmia-associated transcription factor (MiTF), which acts downstream of KITLG-KIT signaling and regulates the expression of key proteins involved in MC differentiation, activity and adhesion (ED Fig. 7g) ^38, 39^. More detailed analysis of the arteriolar unit showed *Kitl* expression at both the inner lining of CD31^+^ endothelial cells and ACTA2^+^ VSMCs (Fig. 7c–e). Unexpectedly, a periarteriolar single-cell layer of fibroblasts also showed *Kitl* promoter activity (Fig. 7c, f and ED Fig. 7h–k). Periarteriolar MCs located very closely to this yet unappreciated subset of dermal fibroblasts (Fig. 7g,h and Supplementary Video 8), and MCs in direct vicinity of this fibroblast layer were mostly of a mature mMCP- 6^high^ phenotype (Fig. 7i). Thus, integrin-mediated adhesion supports MC localization along the arteriolar unit, where several stromal cell types provide a KITLG-rich niche for a mature MC phenotype.

**Fig. 7.**
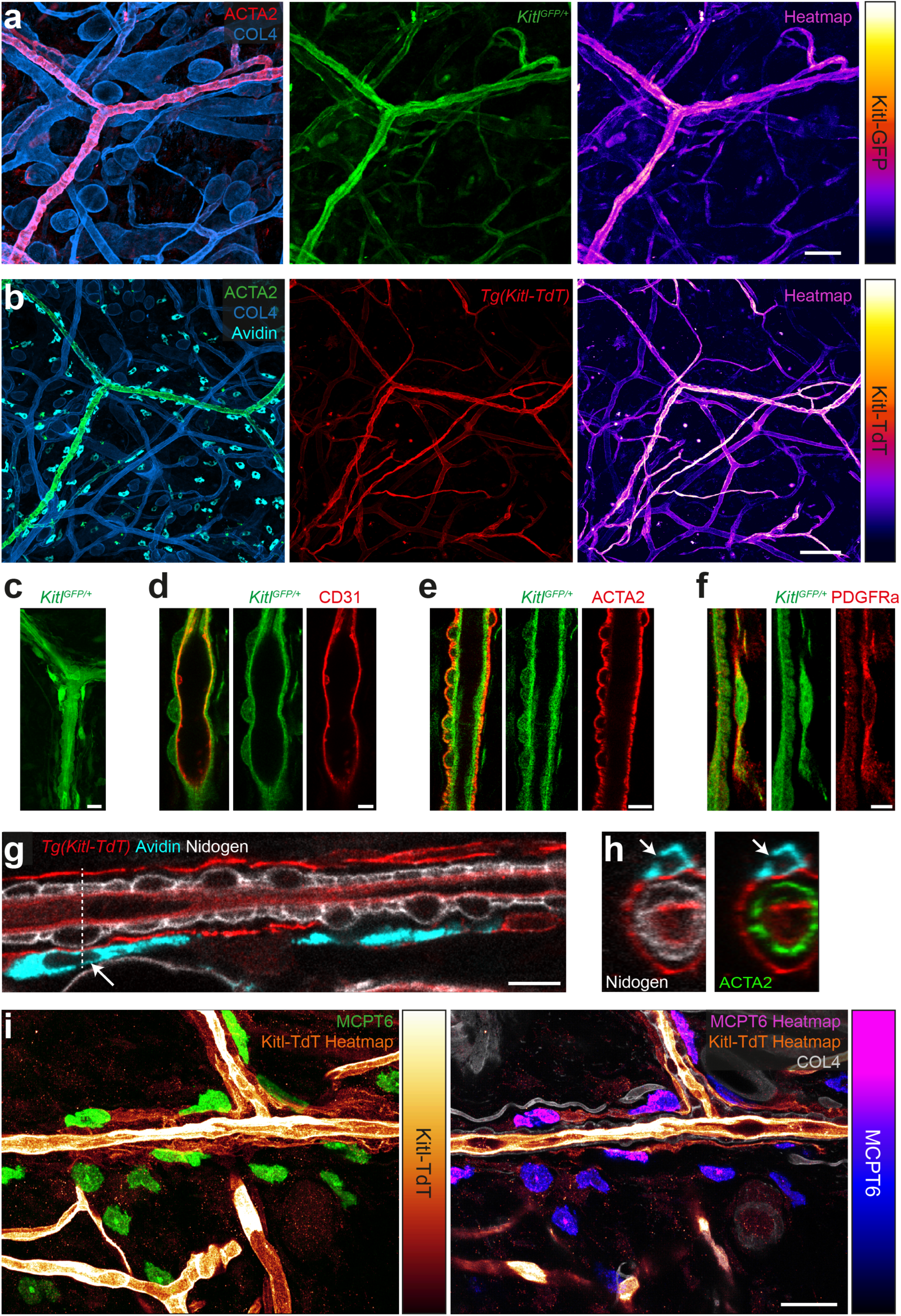
KITLG-expressing stromal cells in the periarteriolar space interact with the mature MC subset. **(a, b)** *In situ* analysis of *Kitl* promoter activity was performed in the ear dermis of two reporter mouse strains. Ear skin whole mount tissues of *Kitl^GFP/+^* (a) and *Tg(Kitl-TdT)* (b) were counterstained for ACTA2-expressing arterioles, COL4-positive basement membranes and MCs (avidin, b only). Representative examples of *n*=5 mice are shown. Endogenous fluorescence intensities are also displayed as heatmaps. **(c–f)** Characterization of GFP- expressing stromal cell types along dermal arterioles of *Kitl^GFP/+^*mice (c). Immunofluorescence stainings were performed against CD31 (endothelial cells) (d), ACTA2 (vascular smooth muscle cells) (e) and PDGFRa (fibroblast) (f). **(g, h)** Periarteriolar positioning of avidin-positive MCs (cyan) and interactions with periarteriolar fibroblasts in *Tg(Kitl-TdT)* mice: (g) longitudinal view of arteriole, (h) cross-section with arteriolar stromal cell layers. White arrows and dashed line indicate the MC in the cross-section area. **(i)** Immunofluorescence analysis of ear skin tissue of *Tg(Kitl-TdT)* mice reveals close proximity and interaction of MCPT6^high^ cells with periarteriolar Kit ligand-expressing fibroblasts. Representative images are volume projection (left) and focal *z*-plane (right). Scale bars: 50 µm (a), 100 µm (b), 15 µm (c), 5 µm (d, f, h), 10 µm (e, g), and 30 µm (i). See also Extended Data Fig. 7 and Supplementary Video 8.

## DISCUSSION

Previous work has shown that the survival and locomotion of amoeboid-like immune cells in interstitial tissues does not require adhesive interactions with the extracellular environment. Adaptive cell shape changes driven by the actomyosin cytoskeleton are sufficient to generate the forces for productive amoeboid leukocyte movement ^6–9, 40^. We here show that MC movement fundamentally differs from this prevailing paradigm widely applied to almost all immune cells: MCs essentially require integrin-mediated force coupling to ECM for their slow interstitial movement, which is crucial for organizing MC distribution in physiological tissue. We recently showed that tissue-resident macrophages, another tissue-resident immune cell type, also require integrins to maintain their mesenchymal shape, control 3D random motility dynamics and optimize tissue surveillance ^41^. However, loss of β1 integrins does not alter their general tissue organization as observed for MCs. Integrin-deficient macrophages still show flexible behavior and productive migration along chemoattractant gradients ^41^, similar to integrin-deficient dendritic cells, neutrophils and T cells, which can move in tissues and 3D gel systems by switching to another mechanistic mode of migration ^3^. In stark contrast, MCs hardly show this plasticity to compensate the loss of integrin-dependent adhesion, which makes MCs special among immune cells and sets them at the outer end of the spectrum of leukocyte migration strategies. It will be interesting for future studies to address why the intracellular actin-driven forces insufficiently support alternative modes of 3D migration in this particular immune cell type. Together, our data place the non-amoeboid movement of WT MCs into the broad category of mesenchymal-like migrating cells despite their hematopoietic origin ^42^. 3D migrating MCs appear to be independent of adherent pressure-driven lobopodial protrusions, as found in highly contractile fibroblasts ^43^. Instead, they form adherent lamellipodia and generate traction by transmitting the retrograde force of actin polymerization to the substrate.

We here rigorously tested the physiological relevance of our findings from artificial 3D gels in native tissue environments and investigated the long-lived pool of connective-tissue type MCs in the adult mouse dermis. These cells critically depend on KITLG as the major growth factor for MC survival and differentiation. Complete loss of KIT receptor signaling in mutant mice interferes with endogenous MC development in most tissues, including the dermis. KITLG haploinsufficiency (*Kitl^GFP/+^* mice) leads to a ∼40% reduction in the number of dermal MCs (data not shown), supporting the general assumption that KITLG concentration determines the size of the dermal MC pool. It has long been known from *in vitro* studies that KITLG induces integrin-mediated adhesion and actin remodeling in MCs and MC-like cell lines ^22^, but it has remained unanswered where this process might play a role *in vivo*. Moreover, the exact cellular sources of KITLG in the mouse dermis have never been characterized in detail ^44^. This is surprising, as critical KITLG-expressing stromal cell niches have been identified for other KIT signaling-dependent cell types. The differentiation of thymocyte progenitors depends on close association to KITLG-expressing vascular endothelial cells and thymic epithelial cells in the cortex of the thymus ^37^. Hair shaft precursors generate a KITLG-providing niche for melanocyte differentiation ^45^. Perisinusoidal stromal cells express KITLG to support extramedullary hematopoiesis of hematopoietic stem cells (HSCs) in the spleen ^46^. Leptin receptor-expressing stromal cells and endothelial cells in the bone marrow are major KITLG sources for the maintenance of HSC and HSC progenitors in the bone marrow ^36, 47^. How KIT and integrin receptor signaling cooperate to organize endogenous mast cell pools has remained unknown.

By studying endogenous dermal MCs in *Tln1^ΔMC^* and *Itgb1^ΔMC^* mice, we make several important findings on the role of integrin receptors for MC tissue homeostasis. First, lack of integrin functionality does not interfere with MC numbers and the general size of the dermal MC pool, which appears to depend entirely on KITLG signaling. This stands in contrast to many other mesenchymal cell types, including fibroblasts and endothelial cells, which require β1 integrin signaling for anchorage-dependent growth and survival ^48, 49^. Thus, adhesive MCs are not just “fibroblast-like” immune cells. Second, talin- and β1 integrin-depletion results in MC cluster formation because of failed MC invasion into the interstitial space. Thus, β1 integrin-dependent haptokinesis is crucial for the homogenous MC distribution in the FN-rich dermal interstitium. Third, β1 integrin receptors anchor MCs to a subset of dermal vessels. Previous studies had identified groups of MCs in close association with several types of blood vessels ^20, 21^. Our analysis of homeostatic skin could not reveal a critical role of integrins for localizing MCs to postcapillary venules and capillaries. However, integrins were absolutely crucial for MC positioning at dermal arterioles. Periarteriolar MC alignment had previously been recognized ^50^, but the physiological function of this spatial correlation has remained unclear. Our results shed new light into this previously unappreciated anatomical structure and identify arterioles as anatomical “hotspots” for KITLG expression in the dermal compartment. Three adjacent stromal cell layers show KITLG expression activity in this region: endothelial cells, VSMCs and a yet uncharacterized type of periarteriolar adventitial fibroblast ^51^. Talin- or β1 Integrin-deficiency restricts most MCs the access and localization to the KITLG-rich periarteriolar space. KITLG can exist both as secreted and membrane-associated form in tissues ^52^. As KITLG is heavily glycosylated and can bind to ECM, a third pool of matrix-associated KITLG is also assumed ^53^. Given that the membrane-bound form could be linked to many physiological and cellular KITLG effects ^37, 54^, we speculate that KITLG may act as haptotactic cue to recruit adhesive MCs into the periarteriolar space and support their localization. However, a chemotactic role, as suggested from *in vitro* studies, cannot be ruled out ^55^. Fourth, localization to the KITLG-rich periarteriolar space promotes a very mature MC phenotype, detected by the upregulated expression of *Mcpt5* and *Mcpt6*, two genes that encode serine proteases abundant in secretory granules of mature MCs ^56^. Based on our observations that mMCP6-expressing periarteriolar MCs appeared in contact with adventitial fibroblasts and VSMC surfaces, direct cell-cell contact with at least one of these stromal cell types might provide KITLG to MCs and induce transcriptional programs for MC maturation ^40^. Integrins do not appear to directly control this MC maturation process, as the few remaining integrin-deficient MCs in direct vicinity to arterioles also show upregulated mMCP6 expression. Thus, the close exposure to high KITLG concentrations appears to be sufficient to promote MC maturation in this tissue niche. How KITLG is exactly presented to MCs in dermal tissue remains unanswered, but future studies involving conditional targeting of KITLG in specific stromal cell types may answer some of these open issues.

In summary, our findings identify MCs as an ECM-anchored cell type that critically depends on substrate adhesion for physiological migration, but not survival. Their slow haptic mode of movement is perfectly adapted to organize long-lasting positioning in tissues with heterogeneous growth factor distribution. Thus, MCs appear unique among immune cells in their migration strategy and “fixed” to a special mode of movement. This stands in stark contrast to the rapid and flexible amoeboid migration of many other immune cell types, whose tissue guidance commonly depends on G-protein coupled receptor signaling ^57^. Furthermore, our results build the basis for future studies assessing MCs as potential mechanosensitive immune cells which probably respond to changing tissue properties under conditions of chronic inflammation, fibrosis and tissue trauma.

## MATERIALS AND METHODS

### Mouse Models

Mice were maintained in a conventional animal facility at the Max Planck Institute of Immunobiology and Epigenetics according to local regulations. Animal breeding and husbandry were performed in accordance with the guidelines provided by the Federation of European Laboratory Animal Science Association and by German authorities and the Regional Council of Freiburg. All mouse strains in this study were without health burden and only used for organ removal after euthanasia by carbon dioxide exposure. *Mcpt5-Cre* ^58^, *Tln1^fl/fl^* ^59^, *Itgb1^fl/fl^* ^60^, *Rosa26^LSL:YFP^* ^61^, *Rosa26^LSL:Tom^* ^62^, *Tg(Ubow)* ^33^, *Tg(Kitl-TdT)* ^37^, *Kitl^GFP^* ^36^, *Tyr^c-2J^* ^63, 64^, *Tg(Myh11-GFP)* ^65^, *Rag2^tm1Fwa^* ^66^, and *Tg(Lifeact-GFP)* ^67^ mouse strains have been described elsewhere. *Mcpt5-Cre Tln1^fl/fl^* mice and crosses with fluorescent reporter lines (*Rosa26^LSL:YFP^*, *Tg(Ubow)*) were on a *Tyr^c-2J/c-2J^*(C57BL/6J-Albino) background, as we initially planned intravital microscopy studies of ear skin in these mice.

### Bone marrow-derived mast cells with connective tissue type characteristics

To obtain mast cells (MCs) with connective tissue type characteristics, we cultured bone marrow-derived MCs (BMMCs) by previously published protocols ^25, 68^. Bone marrow (BM) was isolated by flushing tibiae and femora with cold PBS. Isolated BM cells were maintained at 37 °C and 5% CO_2_ in DMEM (4.5 g/L glucose, Gibco) supplemented with 10% heat-inactivated fetal calf serum (FCS), 10 U/ml penicillin, 10 µg/ml streptomycin, 2 mM L-glutamine, 25 mM HEPES, 1 mM sodium pyruvate, 1x non-essential amino acids (Gibco), 50 µM 2-mercaptoethanol and IL3 (5% supernatant of murine IL3-secreting WEHI-3 cells). To promote BMMC differentiation with connective tissue type characteristics, stem cell factor (SCF; 5% supernatant of murine SCF-secreting CHO transfectants) and IL4 (1 ng/ml, PeproTech) were added to the medium ^25^. IL4 and SCF supplementation to the culture medium enhanced *Mcpt5* promoter activity and thus expression of CRE recombinase in BMMCs generated from *Mcpt5-Cre* mouse strains. A 4-week cultivation of Mcpt5-Cre R26^LSL:YFP^ BMMCs in the presence of IL4 and SCF resulted in 90 to 95% YFP-positive cells in flow cytometry measurements (data not shown). Cells were passaged twice a week and kept at a concentration of 1 to 2.5 × 10^6^ cells/ml as a suspension culture. BMMCs were used for experiments after 5 to 10 weeks of cultivation. Full differentiation into mature MCs was confirmed by expression of lineage-specific c-KIT (rat anti-CD117 Brilliant Violet 421- conjugated, 1:200, Biolegend) and FcεRI (armenian hamster anti-FcεRIα Alexa Fluor 647- conjugated, 1:200, Biolegend) using flow cytometry.

### Peritoneal mast cells

Mouse peritoneal MCs (PMCs) were isolated by peritoneal lavage and maintained at 37°C and 5% CO_2_ in Opti-MEM™ + Glutamax™ (Life Technologies) supplemented with 10% heat-inactivated FCS, 10 U/ml penicillin, 10 µg/ml streptomycin (Gibco) and SCF (5% supernatant of CHO transfectants secreting murine SCF) ^69^. PMCs were passaged twice a week and kept at a concentration of 0.4 to 1 × 10^6^ cells/ml. Experiments with PMCs (5 to 10 weeks of cultivation) were performed after confirming full differentiation into mature MCs using flow cytometric analysis of lineage-specific c-KIT (rat anti-CD117 Brilliant Violet 421-conjugated, 1:200, Biolegend) and FcεRI (armenian hamster anti-FcεRIα Alexa Fluor 647-conjugated, 1:200, Biolegend) expression.

### Dermal mast cells

Dermal MCs (DMCs) were isolated from the ear skin by separating ears into dorsal and ventral halves, mincing tissue and incubating for 75 min at 37°C in RPMI supplemented with 50 U/ml penicillin, 50 µg/ml streptomycin and 0.4 mg/ml Liberase™ (Roche). The tissue digest was stopped by addition of an equal volume of RPMI supplemented with 10% heat-inactivated FCS. To obtain single cell suspensions, digests were subjected to a gentleMACS^TM^ dissociator (Miltenyi; Program C0.1) in C tubes (Miltenyi). After filtration of the suspension through a 70-µm filter (Corning), cells were cultivated for 5 weeks according to the protocol described for PMCs.

### IL3 production by WEHI-3 cell line

WEHI-3 cells were cultured in DMEM (4.5 g/L glucose, 110 mg/L pyruvate; Gibco) supplemented with 10% heat-inactivated FCS, 10 U/ml penicillin, 10 µg/ml streptomycin, 2 mM L-glutamine and 50 µM 2-mercaptoethanol ^70^. Cells were kept at a concentration of 10^5^ cells/ml and medium was exchanged twice a week. For conditioned media production, cells were incubated for 3 to 4 days until the cell concentration reached 1 × 10^6^ cells/ml. Supernatant was collected, centrifuged for 10 min at 350 × g, filtered through a 0.45-µm filter (Nalgene Rapid-Flow sterile disposable filter units with PES membrane) and kept at −20°C for long-term storage.

### SCF production by CHO transfectants

CHO cells were kept at 37°C and 5% CO_2_ in OptiMEM™ + Glutamax™ (Life Technologies) supplemented with 10% heat-inactivated endotoxin-low FCS, 10 U/ml penicillin, 10 µg/ml streptomycin and 30 µM 2-mercaptoethanol. Cells were passaged twice a week and maintained as a confluent culture. For SCF production, cells were seeded at a concentration of 7 × 10^5^ cells per 15 cm cell culture plate (Greiner). When CHO cells reached 70% confluency, supernatant was collected and replaced by fresh medium. CHO supernatant harvest was repeated three to four consecutive days. The supernatant was centrifuged for 5 min at 500 × g, filtered through a 0.2-µm filter (Stericup plastic filter flasks, 0.2 µm) and kept at −20 °C.

### IgE production

Anti-dinitrophenyl (DNP) IgE antibody (clone SPE7) was produced by SPE-7 hybridoma NS1 cells ^71^, which were cultured to a density of 0.5–1 million cells /ml at 37°C and 5% CO_2_ in RPMI supplemented with 10 % FCS, 50 µM 2-mercaptoethanol, and 10 U/ml penicillin. The medium was changed to RPMI supplemented with 1 % FCS, 50 µM 2-mercaptoethanol and 10 U/ml penicillin for two days. Then, the supernatant was collected by centrifugation and sterile filtered. The supernatant was subsequently concentrated using a Sartocon® slice 200 cassette (Sartorius) by the Protein Production Core Facility of the Max Planck Institute of Biochemistry (Martinsried, Germany).

### MEF culture and preparation of 3-D fibroblast-derived fibronectin matrices

Immortalized mouse embryonic fibroblasts (MEFs) were maintained as a confluent culture at 37°C and 5% CO_2_ in DMEM (4.5 g/L glucose, Gibco) supplemented with 10% heat-inactivated FCS, 2 mM L-glutamine, 50 U/ml penicillin, 50 µg/ml streptomycin and 50 µM β- mercaptoethanol and passaged twice a week. Stroma-derived 3D fibronectin (FN) matrices were produced as previously described ^72^. For matrix production, MEFs were seeded at 0.5 × 10^5^ cells/well in 8-well imaging chambers (LabTek^TM^) for live cell imaging or at 1.25 × 10^5^ cells/well in 24-well plates containing round coverslips (12 mm diameter) for static immunofluorescence imaging. Cells were cultured for nine days and fluorescent labeling of matrices for direct visualization during live cell microscopy was achieved by adding 1 µg/ml rhodamine-coupled fibronectin (Cytoskeleton) at day three, five and seven, allowing incorporation of fluorescently labeled fibronectin into the secreted matrices. For static immunofluorescence analysis, the extracted FN matrices were stained with anti-fibronectin antibody (1:200, Sigma) and fluorescently labeled anti-rabbit antibody (Invitrogen).

### Mast cell maturation and integrin expression

To confirm MC maturation and characterize integrin cell surface expression on MCs, flow cytometric analysis was performed. 10^6^ BMMC per sample were used for staining and kept on ice. Antibodies and cells were diluted in PBS supplemented with 2% heat-inactivated FCS (FACS buffer). First, cells were blocked at 4°C for 15 min using anti-mouse CD16/CD32 antibody (1:250, BD Biosciences) before wash. Next, cells were stained for 30 min at 4°C with primary antibodies and washed once with FACS buffer. If primary antibodies were unlabeled, cells were stained with fluorescently labeled secondary antibodies for 30 min. For flow cytometric measurement, MCs were resuspended in FACS buffer and analyzed using a LSRII^TM^ or LSRFortessa^TM^ (BD Biosciences) flow cytometer and FlowJo^TM^ software (BD Bioscience). The following antibodies were used: Alexa Fluor 647-conjugated anti-FcεRI (1:200), Brilliant Violet 421-conjugated anti-cKIT (1:800), PE-conjugated anti-CD29 (1:400), Alexa Fluor 488-conjugated anti-CD49f (1:400), armenian hamster IgG isotype control (1:400) (all BioLegend), PE-conjugate anti-CD11b (1:400), anti-CD18 (1:400), anti-CD49e (1:400), anti-CD51 (1:400), anti-CD61 (1:400), anti-integrin β7 chain (1:400), FITC-conjugated anti-CD41 (1:400) (all BD Biosciences), anti-CD49d (1:400), rat IgG2a kappa isotype control (1:400), rat IgG2b kappa Isotype control (1:400), rat IgG1 kappa isotype control (1:400), rat IgG2a isotype control (1:400) and rat IgG1 kappa isotype control (1:400) (all Thermo Fisher Scientific).

### Determination of talin-1 depletion efficiency

To determine talin-1 depletion efficiencies in BMMC and PMC cultures, intracellular flow cytometry and western blot analysis were performed. For intracellular detection of talin protein expression by flow cytometry, BMMCs were fixed with 2% PFA for 5 min on ice and permeabilized in 0.5% saponin, 0.5% heat-inactivated FCS in PBS (Perm buffer) for 10 min at room temperature (RT) followed by blocking with 2% mouse serum diluted in Perm buffer for 30 min. Cells were washed twice with Perm buffer. Unconjugated anti-pan-talin antibody (1:300, Sigma-Aldrich) was applied in 2% mouse serum-containing Perm buffer for 30 min on ice. Cells were washed twice with Perm buffer and incubated with Cy3-conjugated anti-mouse antibody (1:800, Jackson ImmunoResearch Labs) for 30 minutes on ice, followed by two subsequent washing steps with Perm buffer. Cells were resuspended in FACS buffer for flow cytometric analysis.

For western blot analysis, BMMCs and PMCs were lysed in freshly prepared RIPA buffer (50 mM Tris-HCl, 150 mM NaCl, 0.5% (v:v) NP40, 1% (v:v) Triton^TM^ X-100, 5 mM EGTA, 5 mM EDTA, 1x cOmplete^TM^ protease inhibitor cocktail) for 15 min on ice. Proteins were applied to an 8 to 12% gradient polyacrylamide gel, resolved by SDS-PAGE (BioRad) and transferred onto PVDF membranes (Millipore) via wet blot transfer. Nonspecific binding sites were blocked with 5% skim milk powder in Tris-buffered saline (TBS) containing 0.1% (v:v) Tween-20. Primary antibodies against pan-talin protein (1:1000, Sigma-Aldrich) and actin control (1:2000, Sigma-Aldrich) were incubated overnight in TBS containing 0.1% Tween-20 and 2% bovine serum albumin (BSA; Sigma Aldrich) followed by a 1-hour incubation with HRP-conjugated secondary antibodies (1:5000, Dako) at RT. Upon 5 min incubation with Clarity Western ECL substrate (BioRad), proteins were detected using a ChemiDoc^TM^ Touch Gel Imaging System (Bio-Rad).

### Adhesion assay

To measure MC adhesion to different ECM components and integrin ligands, an absorbance-based adhesion assay was performed. Microtiter plates (Greiner) were coated for 2 h at 37°C in triplicates with fibronectin (10 µg/ml, Sigma-Aldrich), recombinant human ICAM-1 (10 µg/ml, R&D Systems), Matrigel^TM^ (50 µg/ml, Corning) or BSA as control (250 µg/ml) diluted in PBS. Plates were washed twice with PBS and blocked with 3% BSA in PBS for 30 min at 37°C. BMMCs were incubated for 1 h at 37°C in adhesion buffer (phenol red-free RPMI containing 10 mM HEPES, 0.25% BSA, and 2 mM CaCl_2_) or in adhesion buffer supplemented with 1 µg/ml anti-DNP IgE. Cells were washed twice with PBS and seeded at 5 × 10^5^ cells/well in fresh adhesion buffer. Cells were allowed to settle for 10 min at 37°C, 5% CO_2_. IgE-uncoated MCs were stimulated for 30 min with BSA (250 µg/ml), anti-DNP IgE (1 µg/ml) or SCF (100 ng/ml). Anti-DNP IgE-sensitized MCs were stimulated with DNP-HSA (100 ng/ml). After stimulation, plates were centrifuged upside down for 5 min at 60 × g to remove non-adherent cells. After washing once with pre-warmed PBS, plates were centrifuged again upside down. Adherent cells were fixed for 10 min with 4% PFA, followed by a washing step with PBS. Plates were centrifuged 2 min at 420 × g and adherent cells were stained by incubation with crystal violet (5 mg/ml in 2% ethanol) for 10 min at RT. Following two washing steps with tap water, plates were drained upside down and 1% SDS (in H_2_O) was added. Plates were put on an orbital shaker to solubilize the crystal violet dye from lysed cells before absorbance was measured at 570 nm on a Synergy4 plate reader (Bio-Tek).

### Live microscopy of mast cell spreading

To visualize adhesion-dependent MC spreading, live video microscopy was performed with a spinning-disk confocal microscope (Zeiss) equipped with a stage top incubator (TokaiHit) to generate an ambient atmosphere of 37°C and 5% CO_2_. Eight-well imaging chambers (Lab- Tek^TM^) were coated with 15 µg/ml fibronectin overnight at 4°C. After sensitizing BMMCs (non-fluorescent or Lifeact-GFP expressing) with 1 µg/ml anti-DNP IgE for 1 h at 37°C, cells were washed twice in PBS and resuspended in imaging medium (phenol red-free DMEM, supplemented with 10% heat-inactivated FCS, 10 U/ml penicillin, 10 µg/ml streptomycin, 5% IL3 containing WEHI-3 supernatant and 1 mM CaCl_2_) and plated at 5 × 10^4^ cells per well. Microscopy was started immediately after addition of 60 ng/ml DNP-HSA. For differential interference contrast (DIC) time-lapse videos a Plan-Apochromat 40×/ 1.3 Oil DIC Vis-IR M27 objective (Zeiss) was used and images were acquired every 20 s for 1 h. Multiple tiles were recorded to capture a sufficient number of cells. Image tiles were stitched with ZEN blue software. For confocal spinning-disk microscopy time-lapse videos, a Plan-Apochromat 63×/ 1.4 Oil DIC M27 objective (Zeiss) was used and images acquired every 10 s over 8 min.

For the 3D reconstruction of MC adhesion to plate-bound fibronectin molecules, 30 to 50 µm *z*-stacks of individual Lifeact-GFP expressing MCs were acquired with 0.5 µm *z*-step size. Raw imaging data were processed with Imaris (Bitplane) using a Gaussian filter for noise reduction and displayed as 2-dimensional maximum-intensity projections.

MC adhesion to native fibronectin matrices was investigated as follows: Eight-well Lab-Tek^TM^ imaging chambers containing rhodamine-labeled fibronectin matrices were washed twice with PBS and filled with 30 µl imaging medium. BMMCs were resuspended in imaging medium at a concentration of 10^6^ cell/ml and 70 µl cell suspension was added to the matrices for 1 h at 37°C. To stimulate adhesion, 100 µl SCF (final concentration 100 ng/ml) was added to the well. Time-lapse videos were recorded using a Plan-Apochromat 40×/ 1.3 Oil DIC Vis-IR M27 objective (Zeiss) with image acquisition every 30 s over 30 min. 20 µm *z*- stacks with 0.5 µm z-step size were acquired. Raw imaging data were processed with Imaris (Bitplane) using a Gaussian filter for noise reduction and displayed as 2-dimensional maximum-intensity projections.

For confocal spinning-disk microscopy a Cell Observer SD system (Zeiss) consisting of a CSU-X1 confocal scanner unit (Yokogawa) mounted on an AxioObserver Z1 inverted microscope stand and equipped with Evolve back-illuminated EM-CCD camera (Teledyne Photometrics) was used. Depending on the used fluorochromes, images were acquired using laser-line excitation by 488 nm or 561 nm solid-state lasers. For DIC microscopy, a prime BSI scientific CMOS (sCMOS) camera was used.

### MC migration in confined spaces: Under-agarose assay

MC migration behavior in a confined environment was analyzed in an under-agarose assay. The standard protocol ^73^ was modified to allow observation of (a) MC migration over a time course of 14 h, and (b) MC actin flow dynamics.

a) Custom-made imaging chambers (threaded tops of 50 ml centrifuge tubes (Cellstar) fixed in the centre of 3.5 cm diameter petri dishes) were coated with 10 µg/ml fibronectin in PBS overnight at 4°C. Unbound fibronectin was removed by washing the plates twice with PBS. 0.25 g of UltraPure^TM^ agarose (Invitrogen) were dissolved in 10 ml double-distilled water (ddH_2_O) by boiling and mixed with 30 ml HBSS/DMEM solution (1:1 (v:v) HBSS / BMMC medium supplemented with 20% FCS, 5% Il3-containing WEHI-3 supernatant and 5% SCF-containing CHO supernatant) to obtain a final gel concentration of 6.25 mg/ml. To avoid denaturation of the fibronectin layer, agarose gel was cast when the gel mixture was cooled down to a temperature of 40°C. After polymerization, gels were equilibrated for 1 h at 37°C, 5% CO_2_. BMMCs were sensitized with 1 µg/ml anti-DNP IgE for 1 h at 37°C, followed by two washing steps with PBS. 1 µl of the MC suspension (adjusted to 2 × 10^7^ cells/ml in BMMC medium) was injected under the agarose. The outer ring of the imaging chamber was filled with ddH_2_O to avoid dehydration of the gel during the extended imaging time. Cells were incubated for 4 h at 37°C before live-cell microscopy was started using an Axiovert 200 microscope (Zeiss) equipped with a heating chamber set to 37°C and a CO_2_-controller (Zeiss) adjusted to 5% CO_2_. To analyze MC proliferation and migration, time-lapse videos were recorded using a Plan Apochromat 10×/0.25 Ph1 objective (Zeiss). Images were acquired every 10 min for 14 h.
b) To investigate actin flow dynamics of MCs, we used total internal reflection fluorescence (TIRF) microscopy. Custom-made imaging chambers with glass bottom slides (based on 3.5 cm diameter petri dishes with a 15 mm hole drilled into the centre, which was covered with a 18 mm diameter glass cover slip) were coated with 10 µg/ml fibronectin diluted in PBS overnight at 4°C. Control and *Tln1^−/−^* BMMCs were generated from WT *Lifeact-GFP^+/−^* or *Tln1^ΔMC^ Lifeact-GFP^+/−^* mice, respectively. Agarose gels were casted and equilibrated for 1 h at 37°C, 5% CO_2_. Before imaging, anti-DNP IgE-sensitized BMMCs were directly suspended in 60 ng/ml DNP-HSA solution at a concentration of 5 × 10^7^ cells/ml. 1 µl of the cell suspension was injected under the agarose at five injection sites per well and imaging was immediately started. TIRF microscopy was performed using an Eclipse Ti-E inverted microscope (Nikon) equipped with a A1R confocal laser scanning system (Nikon). Images were acquired every 250 ms for 5 to 10min using a Plan Apo TIRF 60× Oil DIC H N2 objective (Zeiss).

For experiments with cytoskeletal inhibitors, 10^6^ anti-DNP IgE-sensitized BMMCs were incubated in 50 µl inhibitor-containing BMMC medium for 1 h at 37°C. Additionally, the agarose and agarose/HBSS solution were also supplemented with the corresponding inhibitor. Following inhibitors were used: Cytochalasin D (1 µM, Merck) and Y-27632 (30 µM, Merck). DMSO and ddH_2_O were used as vehicle control for Cytochalasin D and Y-27632, respectively.

### Mast cell migration in confined spaces: PDMS microchannels

Custom-ordered microchannels (9 µm and 10 µm in width, 10 µm in height; 4D Cell) were treated according to the manufacturer’s protocol. The channels were washed twice with PBS (Gibco) and coated with 100 µg/ml FN in PBS for 1 h at RT. Microchannels were washed three times each, first with PBS and afterwards with BMMC medium. The medium was removed after the last wash step and 10 µl cell suspension (10^7^ cells/ml) were pipetted into each access port. After incubation at 37°C and 5% CO_2_ for 30 min, 2 ml of BMMC medium was added and again incubated at 37°C and 5% CO_2_ for 4 h before imaging was started. Brightfield live cell imaging was performed with an Axiovert 200 microscope (Zeiss) equipped with a heating chamber (Tempcontrol 37-2 digital) set to 37°C and a CO_2_-controller (Zeiss) set to 5% CO_2_. Cells were imaged in a 25- to 30-µm *z*-stack with 2-µm stepping every 10 min for 36 h using a Plan Apochromat 10×/0.25 Ph1 objective (Zeiss). For high-magnification imaging of Lifeact GFP cells in microchannels, a confocal spinning-disk microscope (Zeiss) equipped with a stage top incubator (TokaiHit) was used. Cells were imaged in a 25- to 30- µm *z*-stack with 0.9 µm stepping using a Plan Apochromat 40×/1.3 Oil M27 objective (Zeiss) every minute for 14 min.

### Mast cell seeding of 3D fibrillar space: Matrigel assay

To assess MC migration and seeding in 3D fibrillar matrices, BMMCs were kept in Matrigel™ for up to 72 h. Matrigel™ was used according to the manufacturer’s protocol. BMMCs were sensitized with 1 µg/ml anti-DNP IgE for 1 h at 37°C and 5% CO_2_. After washing twice with PBS, cells were suspended in BMMC culture medium at a concentration of 10^6^ cells/ml. Next, the BMMC suspension was mixed in a 1:1 ratio with BMMC medium containing 20% heat-inactivated FCS, 10% IL3-containing WEHI-3 supernatant and 10% SCF-containing CHO supernatant on ice. 25 µl of this cell suspension was then mixed with 25 µl Matrigel™ on ice to obtain a cell density of 2.5 × 10^4^ cells per 100 µl Matrigel™ solution. For imaging, 10 µl of the gel mixture (2500 cells in a 50% (v/v) Matrigel™) per well were loaded on an µ- slide angiogenesis (Ibidi, cat no. 81506) and kept at RT for 5 min and another 10 min at 37°C and 5% CO_2_ to allow gel polymerization. After that, 50 µl of BMMC medium with 2× concentrated cytokines (10% IL3-containing WEHI-3 and 10% SCF-containing CHO supernatant) was added on top of the gel to provide adequate culture conditions. The prepared samples were pre-incubated for 24 h at 37°C and 5% CO_2_ and imaged afterwards. Live cell imaging was performed using an Axiovert 200 phase-contrast microscope (Zeiss) equipped with a heating chamber and a CO_2_-controller (Zeiss) to generate an ambient atmosphere of 37°C and 5% CO_2_ as described before. Time-lapse videos were recorded up to three days with image acquisition every 10 min and multiple positions to record sufficient cluster formation events using a Plan Apochromat 10×/0.25 Ph1 objective (Zeiss). To correct for a possible optical drift, *z*-stacks with 10 µm step size were acquired. Medium was exchanged every two days. For cluster growth analysis, Matrigel™-embedded cells were incubated at 37°C for seven days, before they were imaged. For experiments in the presence of cytoskeletal inhibitors, 10^6^ anti-DNP IgE-sensitized BMMCs were incubated in 50 µl inhibitor-containing BMMC medium for 1 h at 37°C. Additionally, inhibitors were added into both the Matrigel™ and the BMMC medium that was loaded on top of the solidified Matrigel™. The following inhibitors were used: Blebbistatin (50 µM, Merck), Cytochalasin D (1 µM, Merck) and Y-27632 (30 µM, Merck). DMSO was used as vehicle control for Blebbistatin and Cytochalasin D whereas ddH_2_O was used for Y-27632.

### Ear skin whole mount immunofluorescence analysis

Ears were cut at the base and subsequently split into ventral and dorsal halves. Ventral ear sheets were incubated in 1% PFA for 6 to 8 h at 4°C on a rocker. Ear slices were then transferred to wash and blocking buffer (PBS, 0.25% (v/v) Triton^TM^ X-100, 1% BSA) and incubated overnight by gently shaking at 4°C. After fixation and permeabilization, immunofluorescence staining was performed. During all steps ears were kept at 4°C on a rocker. Each step was followed by three 15 min wash steps with wash buffer. Primary antibodies or directly fluorescent reagents were applied overnight at 4°C. If required, this was followed by a 4 to 6 h incubation at 4°C with fluorescent secondary antibodies. Tissue samples were mounted onto glass slides and covered by glass cover slips (the dermal side facing the cover slip) with Fluoromount-G (SouthernBiotech). Ear explants of Ubow mice were fixed for only 4 h in 1% PFA at 4°C to reduce quenching of the weak CFP fluorescence signal. Samples were otherwise treated according to the described protocol above. The following antibodies were used: anti-ACTA2 Alexa Fluor 405 (1:500, Abcam), anti-ACTA2 Cy3 (1:500, Sigma-Aldrich), anti-CD31 (1:150, BD Pharmingen), anti-collagen IV (1:500, Abcam), anti-ACKR1 Alexa Fluor 488 (1:300, clone 6B7, provided by Aude Thiriot and Ulrich von Andrian, Harvard Medical School, USA) ^74^, anti-desmin (1:300, Abcam), anti-endomucin (1:500, Invitrogen), anti-fibronectin (1:200, Sigma-Aldrich), anti-Lamc1 (1:300, Sigma-Aldrich), anti-Mcpt6 (1:600, R&D Systems), anti-MITF (1:100, Sigma), anti-NG2 (1:200, Sigma-Aldrich), anti-PDGFRa (1:200, Invitrogen) and anti-vimentin (1:300, Abcam). Anti-rabbit Alexa Fluor 405 (1:300, Thermo Fisher Scientific), anti-goat Alexa Fluor 488 (1:500, Invitrogen), anti-rabbit Alexa Fluor 568 (1:500, Thermo Fisher Scientific), anti-rabbit Alexa Fluor 488 (1:500, Invitrogen), anti rat Alexa Fluor 647 and anti-rat Alexa Fluor 405 (both 1:500, Abcam). Avidin was used to visualize dermal MCs: avidin FITC (1:2000, BioLegend) and self-conjugated avidin Alexa Fluor 647 (1:5000; Invitrogen) using an antibody labeling kit (Thermo Fisher Scientific). For signal amplification of endogenous Tomato signal in Tg(*Kitl- TdT*) mice, anti-DsRed (1:1000, Rockland) and anti-rabbit Alexa Fluor 568 (1:500, Invitrogen) were used in some experiments. To amplify the endogenous YFP in *R26^LSL:YFP^* carrying mouse strains, anti-GFP Dylight™ 488 (1:1000, Rockland) was used.

Either *z*-stacks or single focal planes were acquired (indicated in the respective analysis section and/or figure legend). Standard ear whole mount microscopy was performed using an inverted LSM 780 microscope (Zeiss). 3 × 3 tile images were acquired covering 16 to 30 µm of *z*-stack with a step size of 1 or 2 µm using a Plan Apochromat 20×/ 0.8 air objective (Zeiss). Tiled images were stitched during post-processing with ZEN blue software. Images were acquired using excitation through solid-state lasers (UV405; Argon488, DPSS561, HeNe633). The internal photomultiplier tubes and GaAsP detector of the confocal system were used for collecting the emitted fluorescence light.

### Static immunofluorescence analysis of adhesion structures in mast cells

Static immunofluorescence (IF) staining of adhesive structures in adherent MCs were performed on circular glass coverslips (12 mm diameter). Coverslips were coated overnight at 4°C with 15 µg/ml FN diluted in PBS. BMMCs were sensitized with 1 µg/ml anti-DNP IgE for 1.5 h at 37°C and 5% CO_2_. Cells were washed twice with PBS and adjusted to 1.5 × 10^6^ cells/ml in assay medium (BMMC medium supplemented with 5% IL3 containing WEHI-3 supernatant and 2 mM CaCl_2_). Unbound fibronectin was aspirated from the cover slips, which were washed twice with PBS. Cells were plated in 400 µl medium per well and allowed to settle for 10 min at 37°C and 5% CO_2_. Next, 400 µl stimuli solution (recombinant SCF (200 ng/ml, Peprotech) and DNP-HSA (200 ng/ml) was added and cells were incubated for 30 min at 37°C and 5% CO_2_. Floating cells were aspirated and adherent cells were fixed by adding pre-warmed 1% PFA (37°C; diluted in PBS) for 15 min at 37°C. Cells were washed twice with PBS, before PFA was quenched with 0.1 mM glycin for 5 min at RT. Afterwards, the cells were permeabilized with 0.2% Triton^TM^ X-100 in PBS for 5 min at RT and blocked for 30 min with 1% BSA. Primary antibodies were applied 1:200 in 1% BSA overnight at 4°C. Coverslips were washed three times with 1% BSA for 5 min each. Cells were then stained with secondary antibodies (diluted in 1% BSA) for 90 min at RT. After three washing steps for 15 min each, coverslips were rinsed once with ddH_2_O and mounted onto glass slides with antifade mounting medium. The following antibodies were used: Anti-talin (1:200, Sigma-Aldrich), anti-vinculin (1:200, Sigma-Aldrich), anti-CD29 (1:200, BD Bioscience), anti-paxillin (1:200, BD Bioscience) and anti-CD49e (1:200, BD Bioscience). Anti-rat Alexa Fluor 568 (1:500; Invitrogen), anti-mouse Alexa Fluor 647 (1:500, Invitrogen) and anti-mouse Cy3 (1:1000, Jackson ImmunoResearch) were used as secondary antibodies. The actin cytoskeleton was visualized by using phalloidin conjugated to Alexa Fluor 488 (1:2000, Thermo Fisher Scientific). Imaging was performed at a LSM 780 (Zeiss) confocal microscope using a Plan Apochromat 63×/ 1.4 Oil objective (Zeiss). *Z*-stacks were acquired in a range of 7 to 10µm with an step size of 0.51 µm. For deconvolution, cells were acquired starting above and ending under the object of interest, where relevant structures were slightly out of focus with a voxel size of 0.05 µm/voxel in *xy* and 0.51 µm/voxel in *z*. Huygens (SVI) software was used for deconvolution.

To evaluate the formation of adhesive structures in a more native setting, MCs were allowed to adhere to cover slips (15 mm diameter) coated with fibronectin matrices. BMMCs were treated, stimulated and fixed as described in the IF protocol above. After permeabilization and blocking, primary antibodies (diluted in 1% BSA) were applied overnight at 4°C followed by an incubation with secondary antibodies (diluted in 1% BSA) for 1 h at 4°C. The following antibodies were used: anti-fibronectin (1:200, Sigma-Aldrich), anti-CD49e (1:200, BD Bioscience), anti-rabbit 405 (1:500, Thermo Fisher Scientific), and anti-rat Cy3 (1:1000, Jackson ImmunoResearch).

### RNA fluorescence in situ hybridization (RNA-FISH)

To confirm results from RNAseq experiments and analyze differential expression of proteins in ear skin tissue, RNA-FISH by the third-generation in situ hybridization chain reaction (HCR) v3.0 protocol from Molecular Instruments was used according to the manufacturer’s protocol (Molecular Instruments Protocols). Ears from *Mcpt5-Cre^+/−^ R26^LSL:Tom^* mice were dissected following the before described standard protocol and fixed with 4% PFA overnight at 4°C. In an RNAse-free environment, ears were step-wise dehydrated in gradually increasing concentrations of methanol (25%, 50%, 75% and 100%) diluted in PBST (0.1 % Tween 20 in PBS) for 5 min each at RT and kept in 100% methanol overnight at −20°C. Ears were re-hydrated in gradually decreasing concentrations of methanol (100%, 75%, 50% and 25%) for 5 min each at RT before treated with 10 µg/ml proteinase K solution (diluted in PBST) for 2 min at RT. Following a washing step, ears were fixed in 4% PFA for 20 min at RT. For the detection stage, ears were incubated with pre-heated (37°C) hybridization buffer for 5 min at RT and for another 30 min at 37°C for pre-hybridization of the tissues. 250 µl of FISH probes (10 nM) were applied in hybridization buffer and incubated overnight at 37°C. For the amplification stage, ears were incubated in 250 µl amplification buffer containing 30 pmol of fluorescently labeled hairpin 1 and 2 as well as primary antibodies overnight at RT. Following three washing steps with 5x SSCT (5× saline-sodium citrate buffer in distilled water, 0.1% Tween 20), ears were stained with secondary antibodies diluted in 5× SSCT for 4 h at 4°C. Following three washing steps in PBST at RT, ears were mounted onto glass slides using Fluoromount-G (SouthernBiotech). The following HCR probes and antibodies were used: *Tpbs2 (Mcpt6)* (10 probe pairs, Molecular Instruments) and *Cma1* (10 probe pairs, Molecular Instruments), HCR amplifier Alexa Fluor 488 (Molecular Instruments), anti-nidogen (1:300, Merck Millipore), anti-rat Alexa Fluor 405 (1:500, Abcam). Imaging was performed at a LSM 880 (Zeiss) equipped with an Airyscan detector and a Plan Apochromat 20×/0.8 M27 objective (Zeiss). Z-stacks with 0.4 µm step size and 2 x 2 tiles (0.45 x 0.45 mm) were acquired. A 32-channel GaAsP PTM detector of the confocal system was used for collecting the emitted fluorescence light. The FISH signal was detected separately by the integrated Airyscan detector (optimal resolution mode) to improve resolution and signal-to-noise ratio and to avoid channel bleed-through.

### Single-cell RNA-sequencing

#### Cell isolation and FACS

All MCs used for single-cell transcriptomic (scRNAseq) analysis were obtained from 7-week-old *Mcpt5-Cre^+/-^ Tln1^fl/fl^ R26^LSL:YFP^* male mice (*n=3*) and age- and sex-matched control *Mcpt5- Cre^+/-^ Tln1^+/+^ R26^LSL:YFP^*littermate mice (*n=3*). Ears were separated into dorsal and ventral halves, finely minced and incubated for 75 min at 37°C in RPMI supplemented with 5% penicillin-streptomycin and 0.4 mg/ml Liberase™ TM (Roche). Digest was stopped by adding an equal volume stopping solution of RPMI supplemented with 10% FCS to the digestion mixture and keeping cells on ice. Single-cell suspensions were obtained using a gentleMACS^TM^ dissociator (Miltenyi; Program C0.1) and filtering the suspension trough a 70- µm filter (Corning). For an unbiased selection of MCs and to avoid MC activation, we sorted for MC-specific YFP expression, rather than using potentially activating lineage specific markers. After excluding dead cells (DAPI) and doublets, MCs were identified as CD45^+^, Lineage^−^ (CD3e^−^, CD19^−^, CD4^−^, CD8^−^, CD11c^−^, NK1.1^−^), YFP^+^ cells. Cells were sorted into 384-well plates using an Aria Fusion II (BD). The following antibodies were used: BV711- conjugated anti-CD45.2 (1:100, BioLegend), PE-conjugated anti-CD3e (1:100, Invitrogen), anti-CD4 (1:100, Invitrogen), anti-CD8 (1:100, Invitrogen), anti-CD11c (1:100, Biolegend), anti-CD19 (1:100, Invitrogen), anti-NK1.1 (1:100, BioLegend), anti-F4/80 (1:100, Thermo Fisher Scientific). All sorted cells expressed high transcript levels of the MC marker *Cpa3* (Fig. S5B). For scRNAseq analysis of vascular smooth muscle cell (VSMC) and pericytes, CD45^−^ GFP^+^ cells were sorted from ear skin digests of *Myh11^GFP/+^* mice.

#### Single-cell RNA amplification and library preparation

Single-cell RNA sequencing was performed according to the mCEL-Seq2 protocol ^34, 75^. Viable cells were sorted into 384-well plates containing 240 nl primer mix and 1.2 µl PCR encapsulation barrier, Vapour-Lock (QIAGEN) or mineral oil (Sigma-Aldrich). Sorted plates were centrifuged at 2200 × g for 10 minutes at 4°C, snap-frozen in liquid nitrogen and stored at −80°C until they were processed. To convert RNA into cDNA, 160 nl reverse transcription reaction mix and 2.2 µl second-strand reaction mix was used. cDNA from 96 cells was pooled together before clean up and in vitro transcription, generating four libraries from one 384-well plate. 0.8 µl AMPure/RNAClean XP beads (Beckman Coulter) per 1 µl sample were used during all purification steps including library cleanup. Libraries were sequenced on an Illumina HiSeq 2500 and 3000 sequencing system (paired-end multiplexing run, high output mode) at a depth of ∼150,000 to 200,000 reads per cell.

### Quantification of transcript abundance

Paired-end reads were aligned to the transcriptome using bwa (version 0.6.2-r126) with default parameters ^76^. The transcriptome contained all gene models based on the mouse ENCODE VM9 release downloaded from the UCSC genome browser comprising 57,207 isoforms, with 57,114 isoforms mapping to fully annotated chromosomes (1–19, X, Y, M). All isoforms of the same gene were merged to a single gene locus. Subsequently, gene loci with >75% sequence overlap were merged. The right mate of each read pair was mapped to the ensemble of all gene loci and to the set of 92 ERCC spike-ins in the sense direction. Reads mapping to multiple loci were discarded. The left read contains the barcode information: the first six bases corresponded to the unique molecular identifier (UMI) followed by six bases representing the cell-specific barcode. The remainder of the left read contains a polyT stretch. The left read was not used for quantification. For each cell barcode, the number of UMIs per transcript was counted and aggregated across all transcripts derived from the same gene locus. The number of observed UMIs was converted into transcript counts using binomial statistics ^77^.

### Single-cell RNA sequencing data analysis

Clustering and visualization were performed using the RaceID3 algorithm ^34^. Cells expressing >2% of Kcnq1ot1, a potential marker for low-quality cells ^78^, were not considered for the analysis. For normalization, the total transcript counts in each cell were normalized to 1 and multiplied by the minimum total transcript count across all cells that passed the quality control threshold of >1,000 transcripts per cell (Poisson-corrected UMIs ^78^). For the MC data, 1895 cells passed the quality control threshold. For the smooth muscle cell data, 330 cells passed the quality control threshold. RaceID3 was run with the following parameters: mintotal = 1000, minexpr = 5, minnumber = 5. Mitochondrial and ribosomal genes as well as genes with Gm-identifiers were used as input for CGenes.

### Differential gene expression analysis

Differential gene expression analysis between cells and clusters was performed using the diffexpnb function from the RaceID3 package. First, negative binomial distributions reflecting the gene expression variability within each subgroup were inferred on the basis of the background model for the expected transcript count variability computed by RaceID3. Using these distributions, a P value for the observed difference in transcript counts between the two subgroups was calculated and corrected for multiple testing using the Benjamini-Hochberg method as described ^79^. UMAP representation was done as previously published ^80^.

### Mast cell spreading on 2D surfaces

To evaluate MC spreading (PMCs, BMMCs and ear skin MCs), snapshots of indicated timepoints were analyzed with ImageJ (V2.1.0/1.53c). The areas of all cells in a region of interest were manually measured. Mean and standard deviation were calculated using GraphPad Prism.

### Analysis of mast cell migration and actin dynamics in confined spaces

Live cell tracking of MCs in under-agarose assays and PDMS microchannels was performed using the manual tracking option of Imaris 9.1.2 (Bitplane). Cells were randomly chosen and tracked manually through all time frames. The average speed was calculated by dividing the total track length through the total time of imaging in min. Actin dynamics were measured and analyzed via total interference reflection (TIRF) microscopy. Background was subtracted from raw TIRF images using ImageJ (rolling ball algorithm, rolling ball radius 25 pixels) and a FFT bandpass filter was applied (3−40 pixels, no stripe suppression, 5% tolerance of direction). Kymographs were analyzed from filtered image sequences using a custom written Matlab program that allows for assembling of kymographs from arbitrary rectangular selections from leading edges of individual cells (code provided at https://github.com/KMGlaser/Bambach-et-al.git). Actin flow speeds were measured from kymographs by determining the slope of hand-drawn lines that follow prominent actin peaks over time.

### Analysis of mast cell cluster formation in Matrigel™

To analyze differences in the cluster formation between *Tln1^−/−^* and control MCs, the areas of 20 randomly chosen MC clusters after seven days of growth in Matrigel™ were measured using ImageJ (V2.1.0/1.53). To quantify MC cluster growth over three days, images were taken every 20 min. Areas of 20 randomly chosen clusters were manually measured using ImageJ.

### Mast cell morphologies in tissues

To assess MC morphology, YFP-reporter expression in *Tln1^ΔMC^ Rosa26^LSL:YFP^* mice (*n*=3) and control littermates (*n*=4) was used. For MC morphology analysis of *Itgb1^ΔMC^* mice (*n=3*) and control littermates (*n=3*), avidin staining was used to determine the cell shape. Ear whole mounts were acquired with an inverted LSM 780 microscope (Zeiss). A Plan-Apochromat 40×/1.4 Oil DIC M27 objective (Zeiss) was used and 15 to 30 µm *z*-stacks with 1 µm step size were acquired with a zoom of 1.2. For 3D reconstruction and morphometric analysis Imaris 9.1.2 (Bitplane) was used. Morphology of MCs was evaluated by measuring the cellular areas manually in ImageJ (V2.1.0/1.53c) and using the shape descriptor “circularity” = 4π × (area/perimeter^2^).

### Mast cell coverage and proximity to vessels

To evaluate MC proximity to different anatomical structures and to measure the coverage of arterioles by MCs, images of ear skin whole mounts were acquired using a LSM 780 microscope (Zeiss) equipped with a Plan-Apochromat 20× M27 objective (Zeiss). 18 to 25 µm thick *z*-stacks with 1 µm step size were imaged. To capture sufficient numbers of cells, 2 × 2 tile images with 10% overlap were acquired and stitched during post-processing with ZEN blue software. Four imaging field of views (including ACTA2-positive structures) per mouse were analyzed. Mice were between 7 to 10 weeks of age. Periarteriolar alignment analysis was performed using Imaris 9.1.2 (Bitplane). After applying background subtraction and a Gaussian filter to all channels, the surface for endomucin (EMCN)-positive structures was created. Arteriolar surfaces were created after generating a new channel by discarding signal from ACTA2 voxel within endomucin surfaces. For venules, another channel was created by excluding signal from ACTA2 voxel outside endomucin surfaces. To retrieve data on MC proximity to ACTA2-positve arterioles and ACKR1-positive venules, the arteriolar surface was created after generating a new channel by discarding signal from ACTA2 voxel within a previously generated ACKR1 surface. For further details on this image analysis, see also Fig. S3C and D. Tissue-resident MCs were identified using the spot function based on the avidin signal and an estimated diameter of 11 µm. MC alignment to arterioles, venules or capillaries was determined using the Xtension function “find spots close to surface” and setting the threshold to 5 µm.

### Determination of mast cell density in tissues

To determine MC density in ear skin tissue, ear skin whole mounts of *Tln1^ΔMC^* mice (*n*=11) and WT mice (*n*=11) with an age between 7 to 10 weeks were imaged using a LSM 780 microscope (Zeiss) equipped with a Plan-Apochromat 20× M27 objective (Zeiss). 18 to 25 µm thick *z*-stacks with 1 µm step size, 2 × 2 tiled images with 10% overlap and zoom of 1.0 were acquired and stitched during post-processing with ZEN blue software. Four imaging fields of view, displayed as maximum intensity projection, were analyzed per mouse using Imaris 9.1.2 (Bitplane). MCs were identified using the spot function based on the avidin signal. MC density was displayed per analyzed tissue area in in mm^2^.

### Proximity of MCPT6^high^ mast cells to arterioles

To evaluate proximity relationships of MCPT6^high^ expressing MCs to arterioles, surfaces for the pan-MC marker avidin and ACTA2-positive structures were created. The mean intensity of the MCPT6 signal within MCs was used to create MCPT6^high^ and MCPT6^low^ channels. To calculate the proximity of MCPT6^high^ and MCPT6^low^ MCs to arterioles we used the Imaris Xtension tool “find spots close to surface”. Only MCs with a distance between 0-15 µm to arterioles were defined as closely located. Cells at ACTA2-positive structures without characteristic arteriolar morphology (capillaries) were manually excluded from the analysis. For further details on this imaging analysis, see also Fig. S6C and D.

### Mast cell arteriolar coverage

Using Fiji, the length of arteriolar walls and the length of MCs in direct contact to arterioles were manually measured in 2-dimensional maximum-intensity projections of acquired *z*- stacks. Coverage of arterioles by MCs was defined as follows:

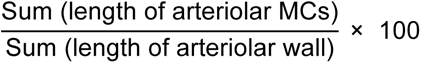

### Ubow analysis

To quantify MC cluster formation in Ubow mice, ear skin whole mounts were imaged using a LSM710 (Zeiss) confocal microscope equipped with a Plan-Apochromat 20× M27 objective (Zeiss). Z-stacks of 25 to 42 µm with 2 µm step size and multiple tiles were acquired. 2 × 2 tiled images (1.2 × 1.2 mm), covering a minimum of 600 MCs, were stitched using ZEN blue software. Statistical analysis and quantification of spatial MC distribution was performed using ClusterQuant2D software ^33, 81^. YFP- and CFP-positive MCs were defined as two distinct classes using the Region-of-interest plugin (ROI manager) of the ClusterQuant2D software. Respective YFP- and CFP-positive MCs were then manually annotated to generate generate cell center coordinate informations. Counterstaining with avidin was used to distinguish MCs from potential auto-fluorescence. Next, 2-dimensional Voronoi meshes were generated based on the cell center coordinates. These diagrams show neighborhood relationships between cells of the same color and allow assessment of spatial distribution of cells in means of diffuse cell spreading or cluster formation (and cluster number/size). These data were used for analysis. For further details on this image analysis, see also Extended Data Fig. 4a. Average of cell numbers per cluster (cluster size) was used for analysis of the clustering index (Extended Data Fig. 4b).

### Quantification of RNA-FISH and MCPT6 expression in arteriolar and interstitial mast cells

Analysis was performed on stitched tile images comprising 25 to 75 MCs using Imaris version 9.5 (Bitplane). First, autofluorescent erythrocytes within the vasculature (using nidogen signal as mask) were excluded. Then, a surface for avidin-positive MCs was created and FISH signal was measured within this mask. Arteriolar aligned and interstitial MCs were manually discriminated. For each class a new surface was created and the according FISH signal was detected using the spot function. Thresholding filters were manually adjusted to detect all FISH signals. The Xtension “split into surfaces” was used to distinguish between FISH spots of individual cells. For quantification, the number of spots (FISH signal) within a cell was divided by the volume of an individual cell.

To evaluate MCPT6 expression in arteriolar and interstitial MCs, images were acquired with a Plan Apochromat 25×/0.8 objective (Zeiss) at a LSM 780 (Zeiss). Z stacks of 25 to 35 µm with 2 µm step size were acquired followed by the stitching of the 2 × 2 filed images using ZEN blue software. After creating a surface for avidin-positive MCs, cells were manually distinguished into arteriolar and interstitial MCs. MCs aligning to ACTA2-positive structures without characteristic arteriolar anatomy were excluded from the analysis. The mean fluorescence intensity per cell of arteriolar and interstitial MCs was quantified.

### Statistical analysis

Analyses were performed using Prism software (GraphPad Software, Inc.Version 8.2.1). If not indicated otherwise, comparisons for two groups were evaluated using a two-tailed unpaired student’s *t* tests after confirming that samples fulfill the criteria of normality. For non-Gaussian distributed data, non-parametric Mann-Whitney U tests were used. Stars indicate significance (\**P*≤0.05, \*\**P*≤0.01, \*\*\**P*≤0.001). NS indicates non-significant difference (*P*>0.05).

## Acknowledgements

We thank R. Fässler, C. Brakebusch, R. Wedlich-Söldner, D. Critchley, S. Monkley, K. Kierdorf for kindly providing mice for this study, J. Friehs for technical help, W. Römer and K. Heger for initial help in this project, members of the MPI Imaging Facility for assistance with imaging, B. Wehrle-Haller for helpful discussions. This work was supported by the Max Planck Society (T.L., D.G.), the Deutsche Forschungsgemeinschaft (DFG, German Research Foundation), Project-IDs 248768892 (T.L.), 431347925 (T.L.) and 322359157 (A.R.), the Else Kröner-Fresenius-Stiftung (2015_A237, M.S.-S.), the Agence nationale de la recherche (ANR-19-CE15-0030, M.B.) and the European Research Council (ERC-2018-COG, ImmuNiche, 818846, D.G.).

## Author contribution

T.L. conceived the project, designed and interpreted experiments. S.K.B., L.K., P.M. and T.L. performed most of the experiments. S.K.B. and T.L. performed the analysis of in vitro and in vivo imaging data. A.G. performed in situ FISH. K.G., M.M. and M.S. assisted with experiments and analysis. R.T. performed TIRF microscopy and analysis. S.K.B. performed statistical analysis. N.A. and D.G. performed scRNA sequencing and data analysis. S.W. and F.K. provided and helped with ClusterQuant2D software. M.S.-S., A.T. and U.v.A. contributed reagents for the study. A.R., M.B. and C.N. contributed mice for the study. M.B. discussed experiments and helped interpret data. T.L. wrote the manuscript.

## Competing Interests

The authors declare no competing financial interest.

**Extended data figures** are available for this paper.

**Supplementary information** is available for this paper.

## Data availability statement

Single-cell RNA-sequencing data have been deposited in Gene Expression Omnibus (GEO) with the accession code GSE205412. The authors confirm that all other data supporting the findings of this study are available within the article and its supplementary materials.

## Code availability statement

Codes for the scRNA-seq data analysis are available upon request through Dominic Grün. ClusterQuant2D code is available through Frederick Klauschen. Kymograph code is provided at: https://github.com/KMGlaser/Bambach-et-al.git

## Notes

### Competing Interest Statement

The authors have declared no competing interest.

