## Supplemental Figures 1-7, Legends to Supplemtal Videos 1-8 for "Slow integrin-dependent migration organizes networks of tissue-resident mast cells"

### Supplemental Information

Extended Data Figures 1-7

Legends to Supplemental Videos 1-8

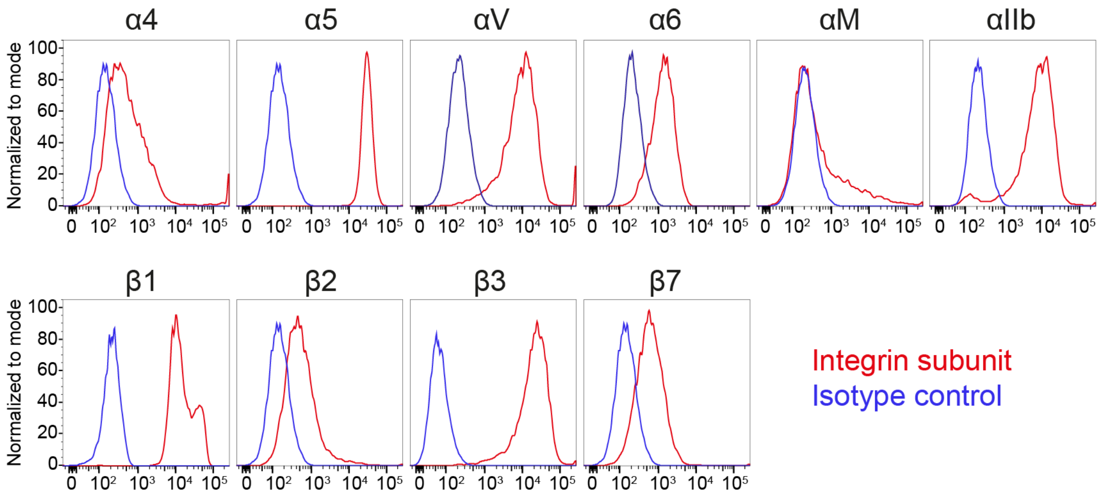


**Extended Data Fig. 1. BMMCs express several members of the integrin family on the cell surface.**
BMMCs that were generated in the presence of IL3, KITLG and IL4 over 6 weeks were analyzed by flow cytometry for the cell surface expression of α and β integrin subunits. Isotype control stainings are displayed in blue, integrin subunit-specific stainings in red.

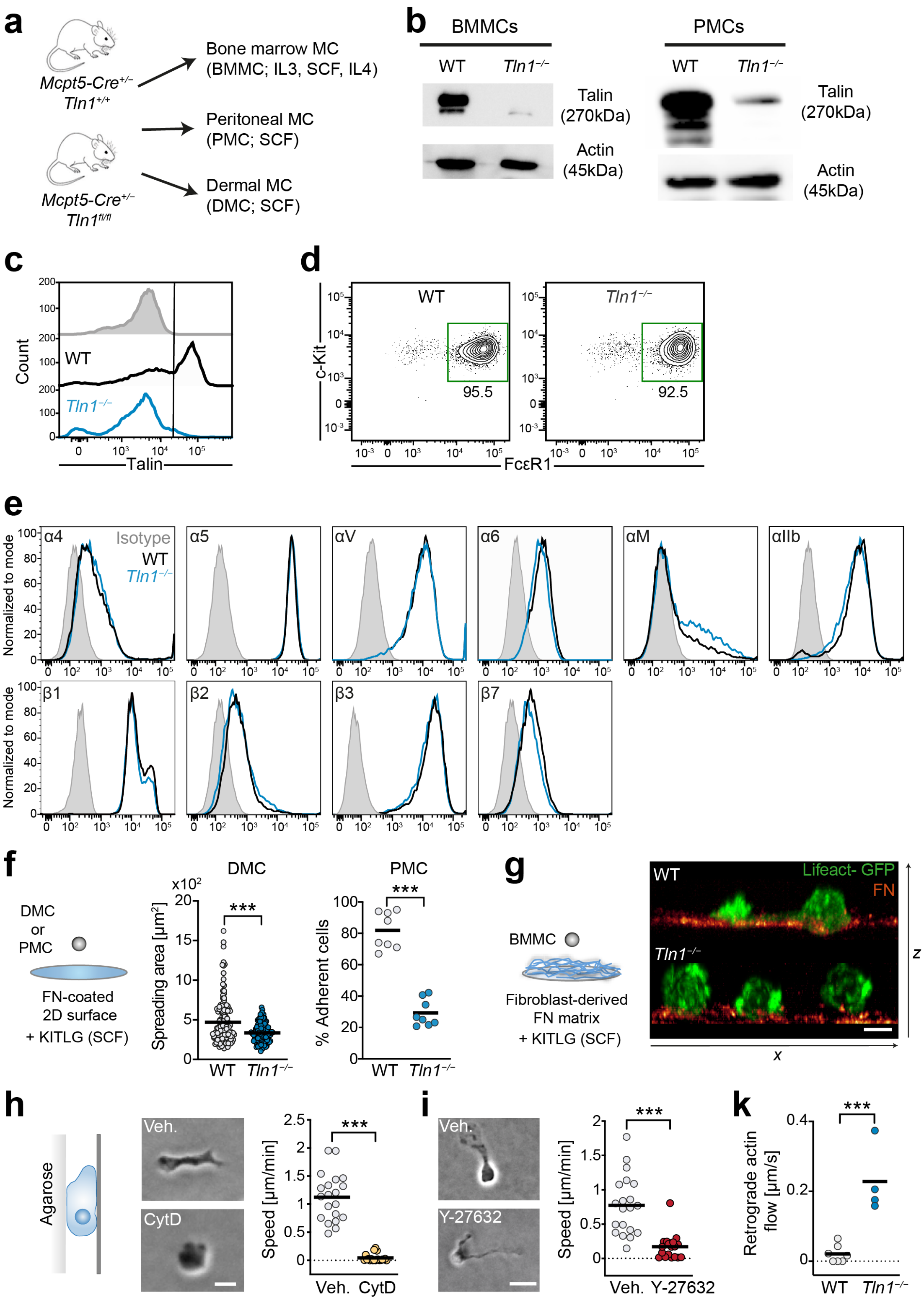


**Extended Data Fig. 2. Functional characterization of *Tln1*$^{-/-}$ MCs and the role of cytoskeletal forces for MC migration.**

**(a)** Scheme for mouse strains and procedures to analyze talin-1 function in several types of MCs, including bone marrow-derived MCs (BMMCs, cultured with additional IL4), peritoneal MCs (PMCs) and dermal MCs (DMCs). **(b, c)** The efficiency of conditional talin-1 knockout was confirmed by immunoblot analysis (b) and flow cytometry (c) by using a pan-talin antibody (recognizing talin-1 and talin-2). Cell lysates were generated from BMMC or PMC cultures of WT and *Mcpt5-Cre*$^{+/-}$ *Tln1*$^{fl/fl}$ mice. Actin was used as loading control (b). Intracellular flow cytometry of BMMCs confirmed the results from the immunoblot. Secondary antibody control is shown on top (grey filled). **(d)** Normal expression of the MC maturation markers c-KIT and FcεR1 in BMMC cultures of WT and *Mcpt5-Cre*$^{+/-}$ *Tln1*$^{fl/fl}$ mice. **(e)** Normal cell surface expression of integrin subunits on *Tln1*$^{-/-}$ BMMCs (in comparison to WT BMMCs, dataset from Extended Data Fig. 1). **(f)** MC spreading of dermal MCs (DMCs) and peritoneal MCs (PMCs) on FN-coated 2D surfaces in response to IgE/DNP-HSA. For DMCs, a comparative analysis of WT and *Tln1*$^{-/-}$ cell spreading area after one hour is displayed. Dots represent values from individual cells from *n*=2 independent experiments for each genotype. Bars display the mean; ****P*<0.001, *t* test. For PMCs, the percentage of adhesive cells was calculated 30 min after stimulation. Dots represent imaging field of views of separate culture wells obtained from *n*=2 independent experiments (with *N*=4 per experiment, i.e. total analysis of more than 100 cells per experiment). ****P*<0.001, *t* test. **(g)** Spinning disk-confocal microscopy of living Lifeact-GFP expressing WT and *Tln1*$^{-/-}$ BMMCs interacting with fibroblast-derived fluorescent FN fibrillar matrix *in vitro*. A snapshot of a confocal *z*-stack is displayed as side view, 2 h after cells were added to the matrix. **(h, i)** The contribution of cytoskeletal forces was analyzed for BMMC migration in the confined space of an under-agarose assay. To interfere with actin polymerization (h) or actomyosin contraction (i), WT cells were treated with cytochalasin D (CytD) or Y-27632, respectively. Representative cell morphologies are displayed and the average cell speed was quantified from one experiment per condition. Dots are values of individual cells (*N*=20 randomly chosen cells per condition). Bars display the mean; ****P*<0.001, *t* test. **(k)** Actin dynamics of Lifeact-GFP expressing WT and *Tln1*$^{-/-}$ BMMCs on FN-coated dishes were recorded with TIRF microscopy and the retrograde actin flow calculated. Each dot represents one cell (*N*=7, WT; *N*=4, *Tln1*$^{-/-}$). Bars display the mean; ***P*<0.01, *U* test. Scale bars: 10 µm (g), 15 µm (h), and 30 µm (i).

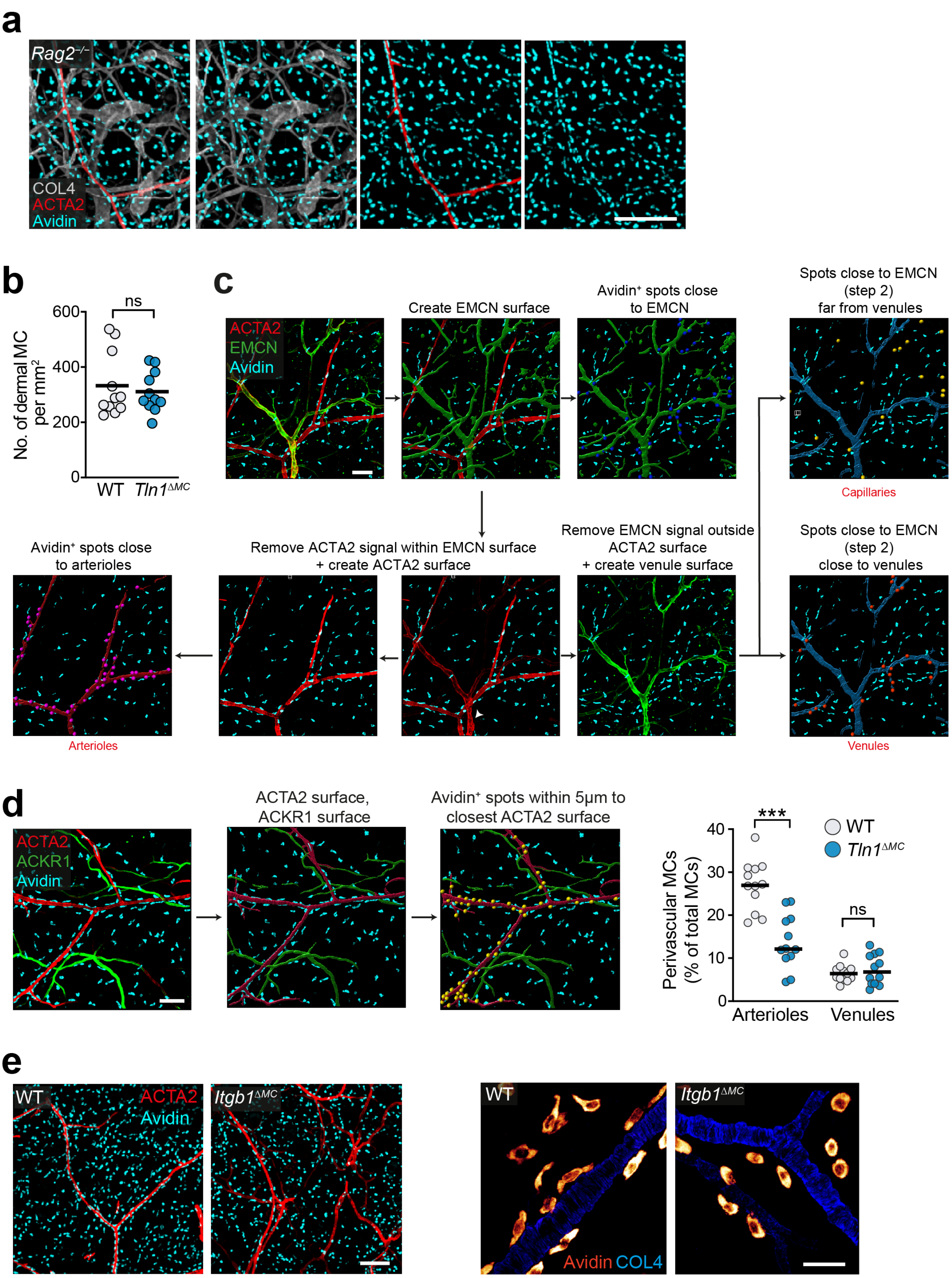


**Extended Data Fig. 3. Analysis of MC network formation and periarteriolar alignment *in vivo*.**
**(a)** Homogenous MC distribution and periarteriolar MC alignment in the dermis of IgE-deficient *Rag2*$^{-/-}$ mice. Representative ear skin whole mount immunofluorescence staining of an 8-week old adult mouse: collagen IV (white), avidin (cyan), ACTA2 (red). **(b)** Histological quantification of MC density in the dermis of ear skin whole mount tissue. Each dot represents the average value of four imaging field of views from one mouse (*n*=11 mice per genotype); *P*>0.05, NS, *t* test. **(c)** Workflow for the post-imaging analysis of MC proximity to arterioles,

venules and capillaries with Imaris software. This analysis was based on immunofluorescence stainings of ear skin whole mount for MCs (avidin), α-smooth muscle actin (ACTA2) and endomucin (EMCN). $ACTA2^+$/$EMCN^-$ vessels were classified as arterioles, $ACTA2^+$/$EMCN^+$ vessels as venules and $ACTA2^-$/$EMCN^+$ vessels as capillaries. This workflow was used for the data analysis in Fig. 3c. **(d)** Workflow for the post-imaging analysis of MC proximity to arterioles, venules and capillaries with Imaris software. This analysis was based on immunofluorescence stainings of ear skin whole mount for MCs (avidin), α-smooth muscle actin (ACTA2) and ACKR1. $ACTA2^+$/$ACKR1^-$ vessels were classified as arterioles and $ACTA2^+$/$ACKR1^+$ vessels as postcapillary venules (see Fig. 3b). MC proximity analysis was comparable to the results obtained with ACTA2/EMCN stainings in Fig. 3c. **(e)** Comparative analysis of dermal MC distribution in adult *Mcpt5-Cre*$^{+/-}$ *Itgb1*$^{fl/fl}$ (*Itgb1*$^{\Delta MC}$) mice and littermate control mice (left, in reference to Fig. 3g). Images of endogenous MC morphologies in native ear dermis are displayed on the right (in reference to the analysis in Fig. 3k). Scale bars: 300 µm (a), 100 µm (c, d), 200 µm (e, left), and 30 µm (e, right).

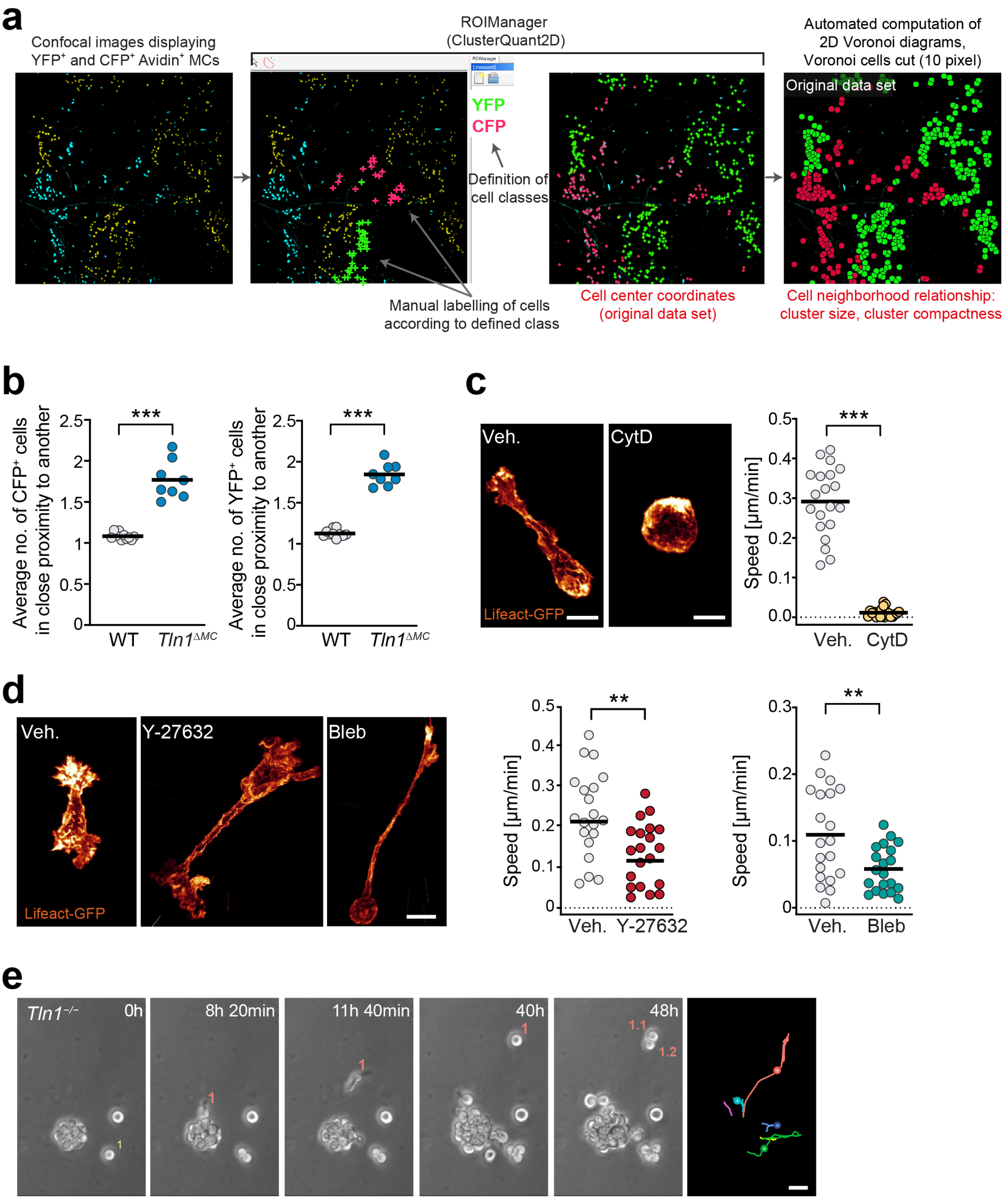


**Extended Data Fig. 4. Analysis of clonal MC invasion and the role of cytoskeletal forces for MC migration in 3D fibrillar environment.**

**(a)** Workflow for the analysis of MC cluster formation with ClusterQuant software. Confocal image stacks from tissues of WT Ubow or *Tln1*$^{\Delta MC}$ Ubow mice (Fig. 4b) form the basis of this analysis. **(b)** Comparative analysis of average YFP$^{+}$ and CFP$^{+}$ cluster size in WT Ubow or *Tln1*$^{\Delta MC}$ Ubow mice. A value of 1 indicates that no cell sits in close proximity to another cell (homogeneous tissue distribution). A value larger than 1 indicates that fractions of cells sit in close proximity to another and form clusters of larger than 1 cell. Dots represent individual imaging fields of view, which come from $n$=3 (WT) and $n$=4 (*Tln1*$^{\Delta MC}$) mice. Bars display the mean; ***$P$<0.001, $t$ test. **(c)** To assess the contribution of actin polymerization for BMMC migration in fibrillar 3D gels, WT cells were treated with cytochalasin D (CytD). Spinning-disk images of Lifeact-GFP expressing WT cells show characteristic cell morphologies in 3D gels. The average cell speed was quantified from one experiment per genotype. Dots are values of individual cells ($N$=20 randomly chosen cells per genotype). Bars display the mean; ***$P$<0.001, $t$ test. **(d)** To assess the contribution of actomyosin contraction for BMMC migration in fibrillar 3D gels, WT cells were treated with either Y-27632 or blebbistatin (Bleb). Spinning-disk images of Lifeact-GFP expressing WT cells show characteristic cell morphologies in 3D gels. Average cell speeds of individual cells were quantified from one experiment. Dots are values of individual cells (*N*=20 randomly chosen cells per experimental condition). Bars display the mean; ***P*<0.01, *t* test. **(e)** Example of a proliferating *Tln1*$^{-/-}$ BMMC cluster in 3D matrigel, which was recorded with live cell microscopy over 48 hours. Rarely, individual cells move away from the proliferating cell cluster and can form a seed for a new proliferating cell cluster (e.g. Cell 1, which moves and then divides to form descendant cells 1.1 and 1.2). Cell tracks over 48 h are shown on the right. Scale bars: 10 μm (c, d), and 30 μm (e).

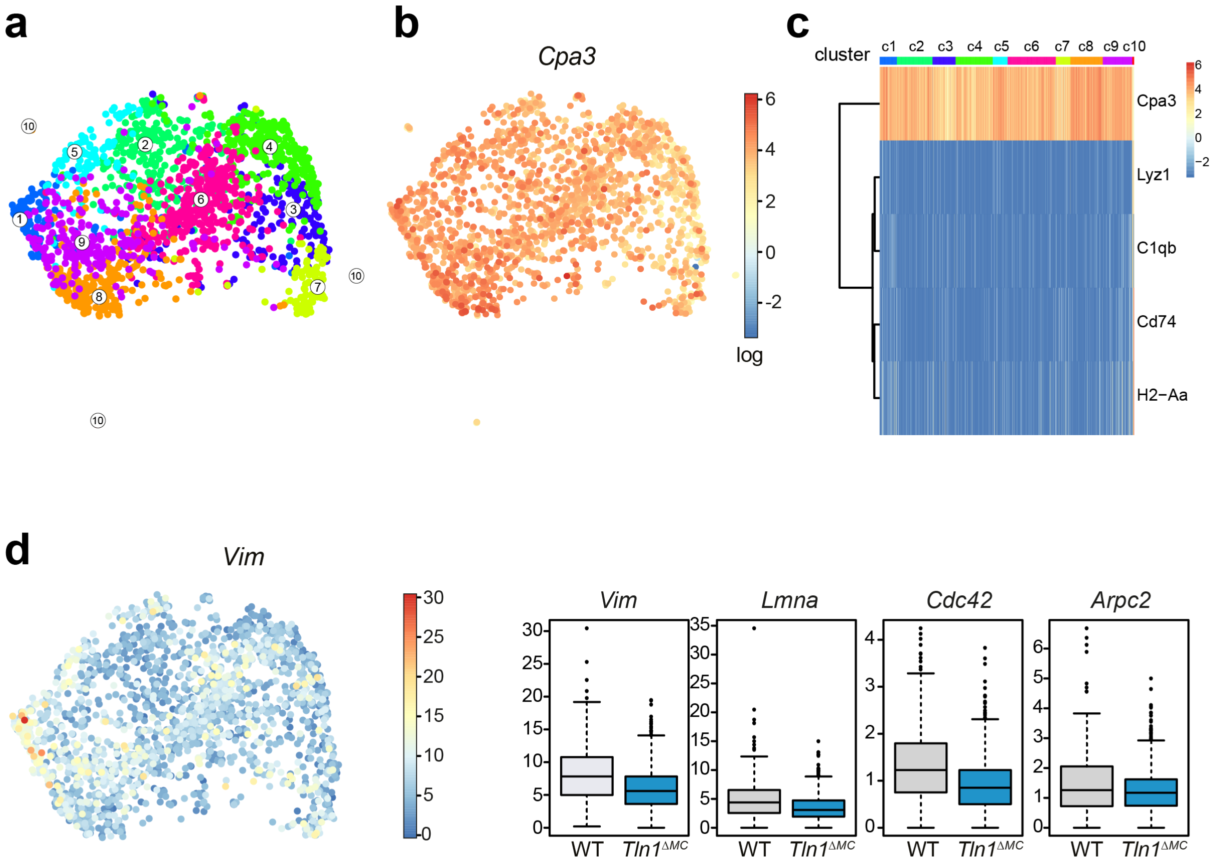


**Extended Data Fig. 5. Single cell RNAseq analysis of dermal MCs.**

**(a)** UMAP of single-cell transcriptomes of sorted CD45$^+$Lin$^-$YFP$^+$ cells (WT and *Tln1*$^{\Delta MC}$ MCs combined) highlighting RaceID3 clusters, displaying clusters 1–10. The outlying cluster 10 was not displayed in the corresponding Fig. 5b, as it carried an additional macrophage gene signature (see Extended Data Fig. 5c). Numbers denote clusters. **(b)** UMAP showing the expression of the MC marker *Cpa3* in clusters 1–10. The color bar indicates log$_2$ normalized expression. **(c)** Heatmap showing the expression of *Cpa3* and markers specifically expressed in cluster 10. The color bar at the top of the heatmap indicates the RaceID3 clusters. Scale bar, log$_2$ normalized expression. **(d)** UMAP showing the expression of *Vim* (left). The color bar indicates normalized expression or transcript counts. Box plots show analysis of clusters 1, 8, 9 and the expression of the cytoskeletal elements *Vim*, *Lmna, Cdc42, and Arpc2* in WT and *Tln1*$^{\Delta MC}$ cells (right). The expression of the genes was significantly downregulated in MCs from *Tln1*$^{\Delta MC}$ relative to WT mice. For all comparisons: Benjamini-Hochberg corrected *P*<0.05 (see Methods).

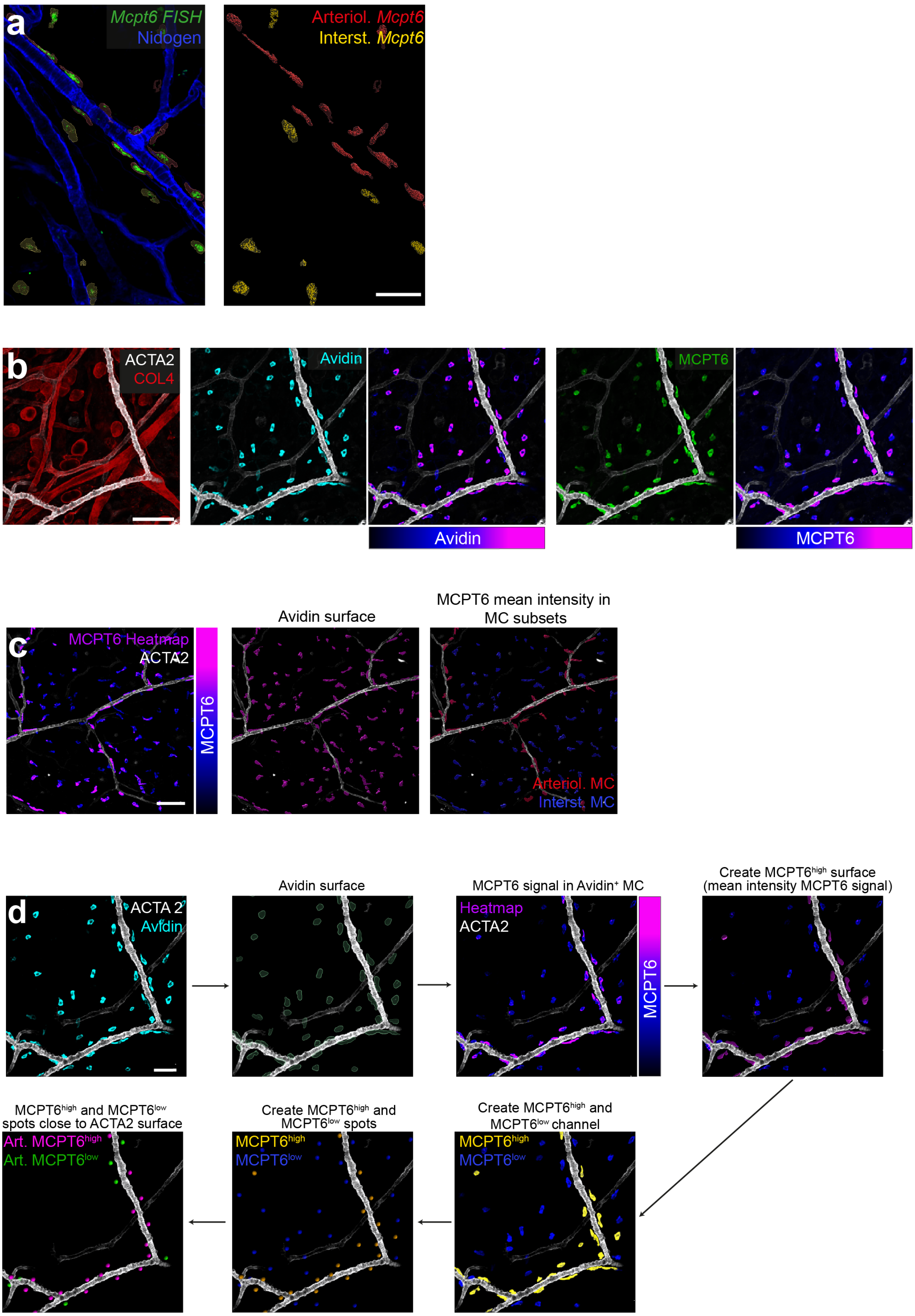


**Extended Data Fig. 6. Analysis of the periarteriolar MC phenotype *in situ*.**
**(a)** Representative visualization of *Mcpt6* mRNA using fluorescence in situ hybridization (FISH) in ear skin whole mount tissue of adult WT mice (left) and post-imaging analysis of *Mcpt6* expression in periarteriolar versus interstitial MCs (right). These data refer to the quantification in Fig. 6c, d. **(b)** Differential expression of MCPT6 in

periarteriolar versus interstitial MCs. Ear skin whole mount tissue of an adult WT mouse is counter-stained for the pan-MC marker avidin (cyan), MCPT6 (green), COL4-expressing basement membranes (red) and ACTA2-positive arterioles (white). Avidin and MCPT6 fluorescence intensities are also displayed as heatmap. **(c, d)** Imaris software-based workflows for the post-imaging analysis of MCPT6 mean fluorescence intensities in MC subsets (d) and calculation of the percentage of Mcpt6$^{high}$ MCs of all periarteriolar MCs in WT and *Tln1*$^{\Delta MC}$ mice (e). Scale bars: 50 µm (a, d), and 100 µm (b, c).

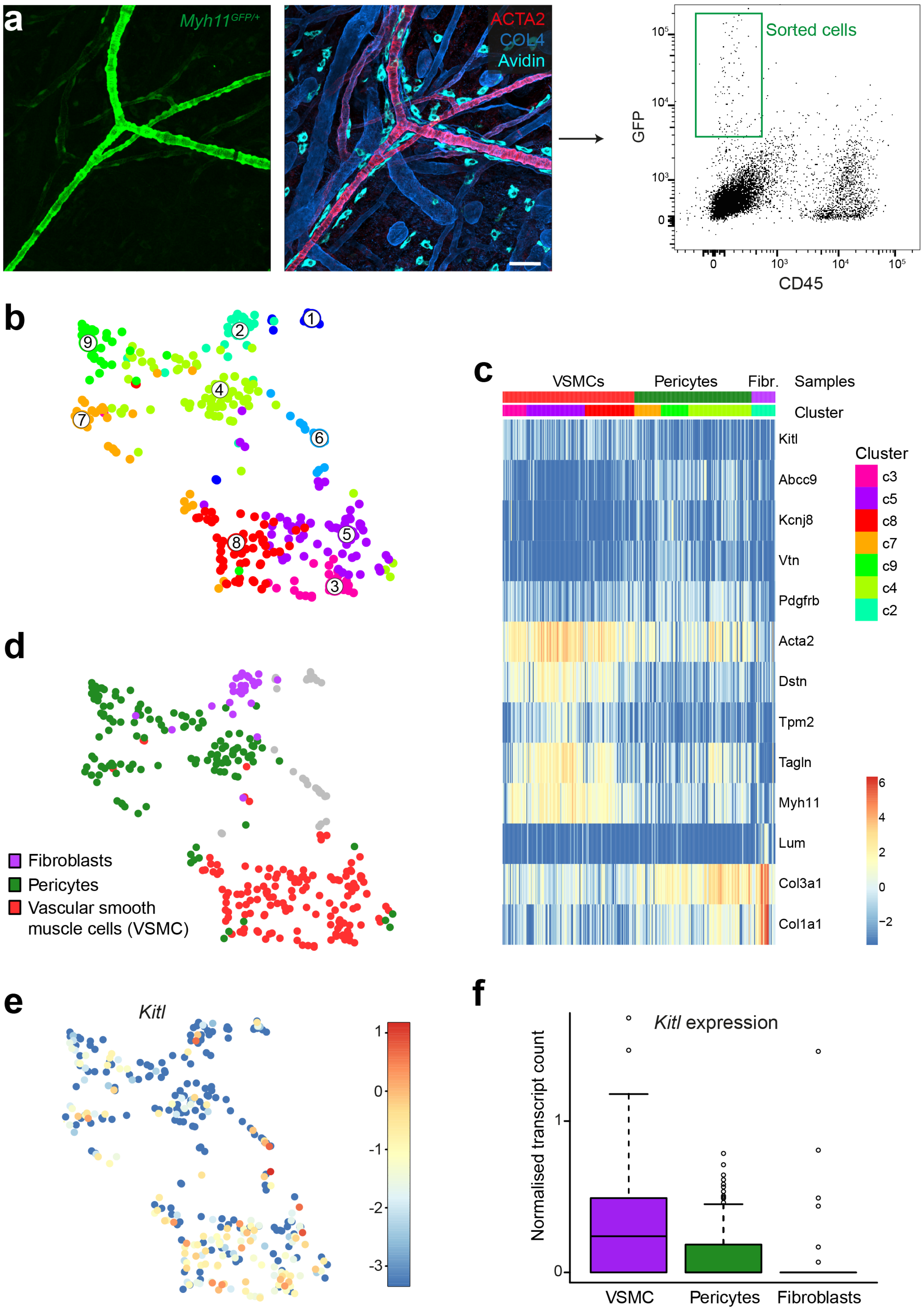

a
Myh11GFP/+
ACTA2
COL4
Avidin
Sorted cells
GFP
CD45
b
c
VSMCs
Pericytes
Fibr.
Samples
Cluster
Kitl
Abcc9
Kcnj8
Vtn
Pdgfrb
Acta2
Dstn
Tpm2
Tagln
Myh11
Lum
Col3a1
Col1a1
c3
c5
c8
c7
c9
c4
c2
d
Fibroblasts
Pericytes
Vascular smooth muscle cells (VSMC)
e
Kitl
f
Kitl expression
Normalised transcript count
VSMC
Pericytes
Fibroblasts

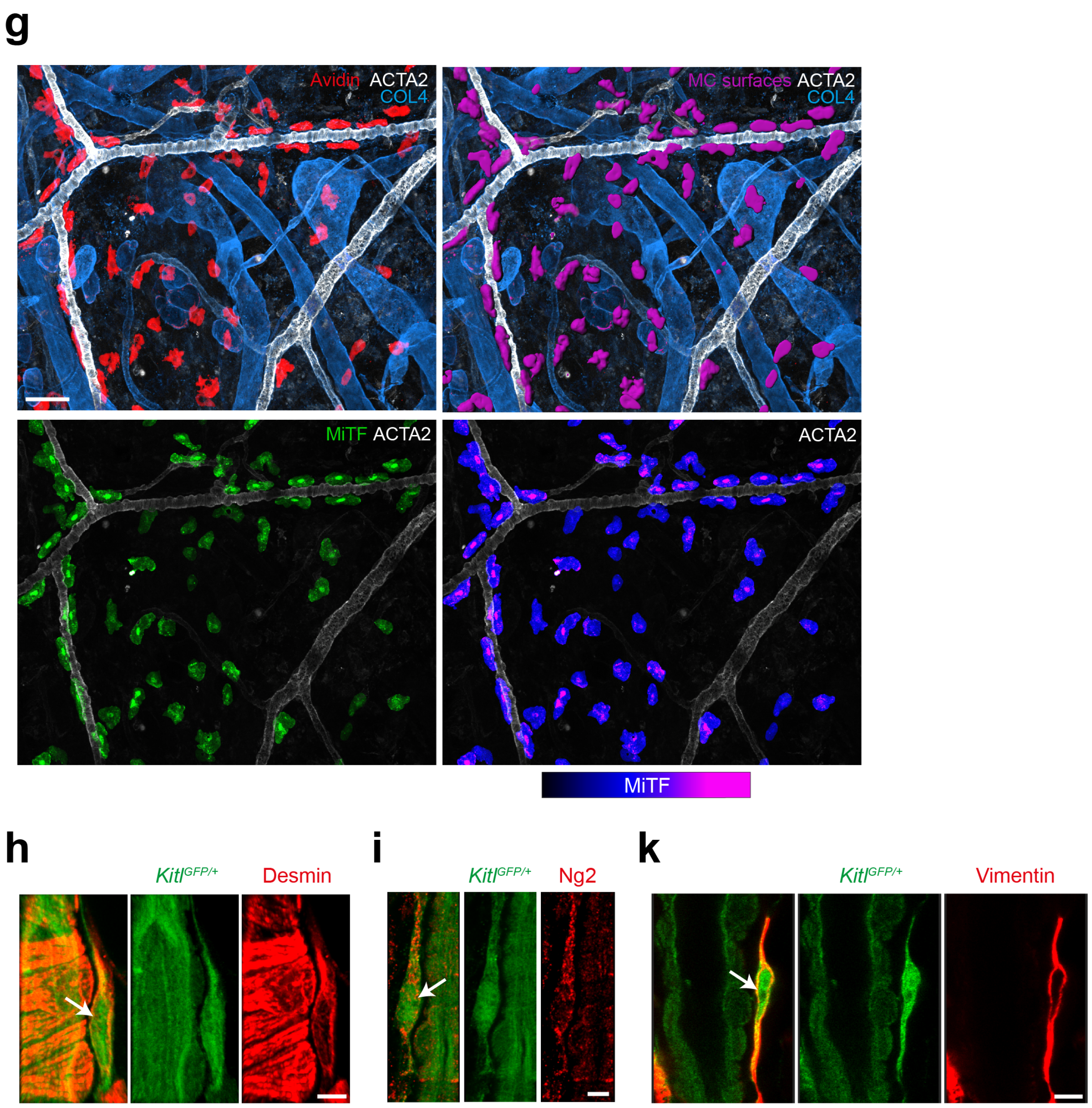


**Extended Data Fig. 7. Analysis of periarteriolar stromal cells with scRNAseq and *in situ* immunofluorescence analysis.**

**(a)** Experimental strategy and work flow for scRNAseq analysis of perivascular stromal cells with focus on Myh11-expressing vascular smooth muscle cells (VSMCs). Myh11-GFP transgenic mice carry a strong genetic label for VSMC and very weak GFP-label for pericytes. Ear skin tissue of a Myh11-GFP mouse was digested, and GFP-fluorescent cells sorted in preparation for scRNAseq analysis. **(b)** UMAP highlighting RaceID3 clusters. Numbers denote clusters. **(c)** Heatmap showing the expression of marker genes for each cell type compartment. Color bars indicate cell type and RaceID3 clusters. Three stromal cell types (VSMC, pericyte, fibroblast) were retrieved after the sort. Scale bar, $\log_2$ normalized expression. **(d)** UMAP highlighting the annotated stromal cell types. **(e)** UMAP showing the expression of *Kitl*. Color bar indicates $\log_2$ normalized expression or transcript count. **(f)** Box plots showing the normalized expression of *Kitl* in VSMCs, pericytes, and fibroblasts. **(g)** Immunofluoresence staining analysis of nuclear MiTF signal in dermal mast cells of ear skin whole mount tissue. MC surfaces were generated based on avidin staining. Only MiTF signal inside MC surfaces is displayed as single color or as heatmap. Many of the periarteriolar positioned MCs show strong nuclear MiTF signals. **(h–k)** GFP-expressing fibroblasts that reside along dermal arterioles in *Kitl*$^{GFP/+}$ mice were phenotypically characterized by immunofluorescence stainings against desmin (h), chondroitin sulfate proteoglycan (Ng2) (i) and vimentin (k). Scale bars: 50 µm (a, g), and 5 µm (h–k).

### Legends to Supplemental Videos

**Supplementary Video 1: Integrin-dependent mast cell spreading on fibronectin-coated 2D surfaces (related to Fig. 2)**
Mast cells were cultivated from $Tln1^{\Delta MC}$ $Lifeact\text{-}GFP^{+/-}$ and WT $Lifeact\text{-}GFP^{+/-}$ mice, coated with anti-DNP IgE and applied to fibronectin-coated 2D surfaces. DNP-HSA was added to induce FcεRI-mediated mast cell spreading. Spinning-disk confocal microscopy of cellular actin revealed that WT mast cells adhered, spread and started to migrate, whereas $Tln1^{-/-}$ mast cells did not adhere and spread. This video relates to Fig. 2c.

**Supplementary Video 2: Integrin-dependent mast cell motility on fibroblast-derived fibrillar fibronectin matrix (related to Fig. 2)**
Mast cells were cultivated from $Tln1^{\Delta MC}$ $Lifeact\text{-}GFP^{+/-}$ and WT $Lifeact\text{-}GFP^{+/-}$ mice and applied to rhodamine-labeled fibronectin matrices in the presence of SCF. Video sequences are displayed on top, static side views are displayed on the bottom. Spinning-disk confocal microscopy revealed that WT mast cells integrate, interact and move into the matrix. In contrast, $Tln1^{-/-}$ mast cells fail to invade the matrix and float on top the fibrillar fibronectin layer. This video relates to Extended Data Fig. 2g.

**Supplementary Video 3: Integrin-dependent mast cell migration in confined space (under-agarose assay) (related to Fig. 2)**
Mast cells were cultivated from $Tln1^{\Delta MC}$ and littermate control mice and applied to an under-agarose assay. Mast cell movement occurred in the confined space between the agarose layer and the fibronectin-coated cell culture plastic underneath. Phase contrast brightfield microscopy over 14 hours revealed random movement and lamellopodia formation of WT mast cells, $Tln1^{-/-}$ mast hardly moved and remained largely immotile. This video relates to Fig. 2f–h.

**Supplementary Video 4: Lack of high-affinity integrins increases the retrograde actin flow in mast cells (related to Fig. 2)**
Lifeact-GFP expressing mast cells were confined between an agarose layer and a fibronectin-coated cell culture plastic, and lamellopodial actin dynamics recorded. Total internal reflection (TIRF) microscopy revealed protrusive lamellopodia movement without measurable retrograde actin flow in WT mast cells, whereas $Tln1^{-/-}$ mast cells displayed a clear increase in retrograde actin flow. This video relates to Extended Data Fig. 2k.

**Supplementary Video 5: Integrin-dependent mast cell migration in confined space (PDMS microchannels) (related to Fig. 2)**
Mast cells were cultivated from $Tln1^{\Delta MC}$ and littermate control mice and applied to the confined space of 10 µm × 10 µm PDMS microchannels. A representative selection of 10 channels per genotype is displayed. Phase contrast brightfield microscopy over 100 minutes revealed clear movement of WT mast cells in these confined spaces. In contrast, $Tln1^{-/-}$ mast cells, which had managed to invade the microchannel, did not show active crawling behavior during this time frame. This video relates to Fig. 2i–m.

**Supplementary Video 6: Mast cell seeding in 3D fibrillar space requires integrin-dependent migration (related to Fig. 4)**
Mast cells were cultivated from $Tln1^{\Delta MC}$ and littermate control mice, applied to 3D matrigels in the presence of SCF and observed over 60 hours by phase contrast brightfield microscopy. The movement of WT mast cells is displayed first and includes fate tracking of two cell clones and corresponding descendant cells. This video shows how proliferation and integrin-dependent migration shape the formation of a cellular network with heterogenous cell distribution. The second part of the video shows how the majority of $Tln1^{-/-}$ mast cells are unable to move away from each other after cell division, which results in the growth of mast cell clusters. This video relates to Fig. 4g–i.

**Supplementary Video 7: Rare example of an individual mast cell escaping a $Tln1^{-/-}$ proliferation cluster and forming a new seed for another cluster (related to Fig. 4)**
$Tln1^{-/-}$ mast cells were applied to 3D matrigels in the presence of SCF and observed over 9 hours by phase contrast brightfield microscopy. The majority of $Tln1^{-/-}$ mast cells are proliferating, but these cells are unable to move away from each other after cell division, which results in the formation of mast cell clusters. This video highlights a rare example of an escapee mast cell, which moves away from the cell cluster and forms a new seed for a new cell cluster. This video relates to Extended Data Fig. 4e.

**Supplementary Video 8: Positioning of mast cells in the SCF-rich periarteriolar niche with direct contact to SCF-expressing periarteriolar fibroblasts (related to Fig. 7)**

This video shows an animated view of SCF expression in the arteriolar tissue niche. Ear skin explants of *Tg(Kitl-TdT)* mice were fixed and counterstained for avidin (mast cell, cyan), α-smooth muscle actin (vascular smooth muscle cells, VSMC) and collagen IV (basement membrane). Confocal fluorescence microscopy and 3D reconstruction of the confocal image stacks reveal a direct contact of periarteriolar mast cells with SCF-expressing fibroblasts, which form an outer cellular layer in the periarteriolar space. This video relates to Fig. 7g, h.
